# Hippocampal place code plasticity in CA1 requires postsynaptic membrane fusion

**DOI:** 10.1101/2023.11.20.567978

**Authors:** Mark H. Plitt, Konstantin Kaganovsky, Thomas C. Südhof, Lisa M. Giocomo

## Abstract

Rapid delivery of glutamate receptors to the postsynaptic membrane via vesicle fusion is a central component of synaptic plasticity. However, it is unknown how this process supports specific neural computations during behavior. To bridge this gap, we combined conditional genetic deletion of a component of the postsynaptic membrane fusion machinery, Syntaxin3 (Stx3), in hippocampal CA1 neurons of mice with population *in vivo* calcium imaging. This approach revealed that Stx3 is necessary for forming the neural dynamics that support novelty processing, spatial reward memory and offline memory consolidation. In contrast, CA1 Stx3 was dispensable for maintaining aspects of the neural code that exist presynaptic to CA1 such as representations of context and space. Thus, manipulating postsynaptic membrane fusion identified computations that specifically require synaptic restructuring via membrane trafficking in CA1 and distinguished them from neural representation that could be inherited from upstream brain regions or learned through other mechanisms.

## Introduction

Learning occurs through experience-dependent changes to neural circuits, thus altering how the nervous system processes future information. Though neurons can accomplish these circuit changes in a myriad of ways, one well-characterized mechanism utilized by mammalian neurons is to increase the number of AMPA-type glutamate receptors (AMPARs) at the postsynaptic density^1–4^. This process results in a lasting increase in the strength of excitatory synapses referred to as long term potentiation (LTP)^5^. These additional AMPARs, which typically contain the GluA1 subunit, are immediately supplied from a pool of extrasynaptic diffusing AMPARs in the postsynaptic membrane^3,6^ or by local exocytosis of AMPAR-containing vesicles^7^. According to one model, LTP-inducing stimuli cause diffusing GluA1(+) AMPARs to be trapped and stabilized at the synapse by interactions with scaffolding elements^8–11^, with exocytosis of additional AMPARs from recycling endosomes being necessary for replenishing or regulating this extrasynaptic poo_l_^12,13^. According to an alternative model, AMPARs necessary for LTP are directly inserted into the postsynaptic specializations. Independent of which model is correct, AMPARs as well as other components of postsynaptic specializations (e.g. NMDA receptors and kainate receptors) must reach the synaptic membrane via regulated exocytosis. Importantly, exocytosis of GluA1(+) AMPARs in response to LTP-inducing stimuli requires a molecularly distinct membrane fusion complex from the membrane fusion machinery used for basal and homeostatic regulation of surface AMPARs^14–16^. Although this AMPAR mobilization framework is a dominant model for synaptic learning, it remains controversial whether LTP-specific membrane fusion proteins are needed for certain forms of synaptic potentiation^17–19^. In particular, it is unclear if activity-dependent membrane fusion is necessary for generating *in vivo* neural dynamics that support learning.

The hippocampal circuit is a powerful model system for studying the role of regulated membrane traffic in driving plasticity of neural codes during behavior. Principal excitatory cells in all hippocampal subfields (e.g. CA1 & CA3) contain “place cells”, a functionally defined cell class that fires action potentials specifically when an animal is in one or several restricted spatial locations^20–23^. Within an environment, place cells tile space via their place fields to form a “cognitive map” of the environment^24^. These firing patterns form rapidly when an animal enters a novel environment and are largely reactivated when an animal reenters the same environment^25–34^. Place cells can also make lasting changes to their firing patterns to incorporate new experiences in an environment^35–37^. Due to the redundancy of spatial representations across hippocampal subfields, it remains a challenge to disentangle the roles of synaptic plasticity and inherited neural dynamics in shaping place cell properties. Manipulating activity-dependent membrane traffic provides a potential means to reduce synaptic potentiation while leaving baseline synaptic integration unaffected.

In addition to gradual changes in hippocampal coding that may occur during behavior, plasticity of CA1 pyramidal cell coding is associated with at least two types of discrete events. First, hippocampal CA1 pyramidal neurons are capable of a rapid form of plasticity called Behavioral Timescale Synaptic Plasticity (BTSP), in which new place cell representations can form after a single sustained dendritic depolarization called a plateau potential^38–44^. Second, sequences of place cells that were active during exploration are stabilized through plasticity during offline reactivation events referred to as sharp-wave ripples (SWRs) - high frequency local field potential (LFP) events in which the same sequences of place cells are reactivated at highly compressed speeds^45–47^. Though the molecular underpinnings of BTSP and SWR-induced plasticity are a topic of increasing interest, the mechanisms for expressing either form of plasticity remain a mystery^38,48–52^.

In this work, we make a direct causal link between postsynaptic membrane traffic and *in vivo* CA1 computations. We performed conditional genetic deletion of Syntaxin-3 (Stx3, Fig. 1A), a necessary component of the LTP-specific postsynaptic membrane fusion complex^14,15^ and other membrane trafficking reactions, in dorsal CA1 neurons of mice before imaging the calcium activity of the same neurons during behavior using two photon microscopy. We found that CA1 Stx3 is not necessary for forming spatially stable place cell representations (i.e. place codes). These place codes can be inherited from upstream synaptic partners or learned through other forms of plasticity. On the other hand, CA1 Stx3 is necessary for novelty processing and spatial reward memory, hippocampal computations that are unlikely to be inherited from upstream synaptic partners. We provide evidence that CA1 Stx3 is necessary for the expression of BTSP-like events in novel environments, implicating rapid exocytosis in this form of plasticity. Lastly, we find that CA1 Stx3 is necessary for offline stabilization of place codes. Collectively, these findings point to a crucial role for activity-dependent exocytosis in sculpting numerous CA1 computations, but they also raise the possibility that synaptic plasticity may not be necessary for maintaining neural dynamics present in upstream inputs under certain conditions.

**Figure 1.**
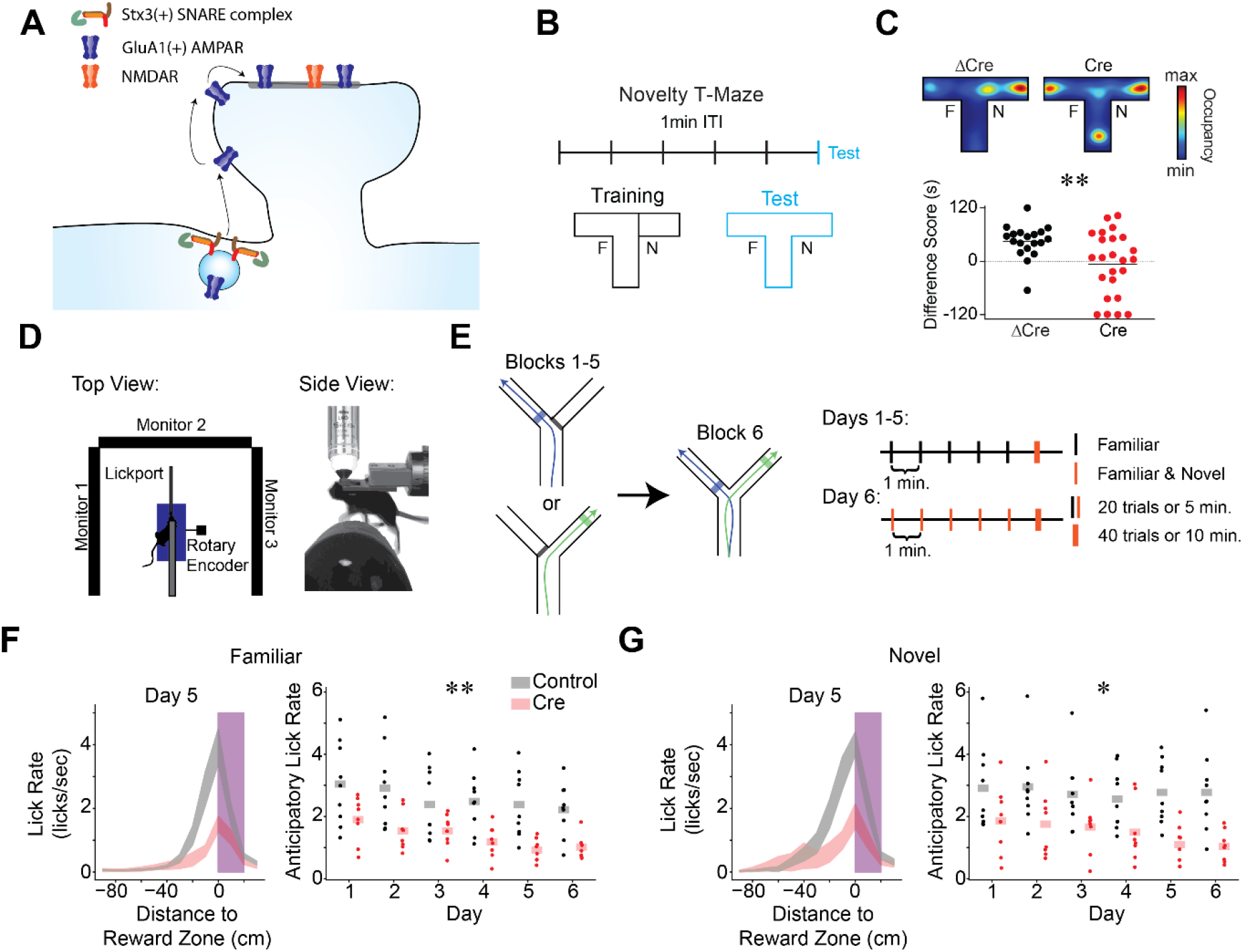
Stx3-dependent postsynaptic membrane fusion in CA1 is critical for novelty preference and reward anticipation. A) Model of Stx3-dependent AMPA receptor trafficking. Stx3(+) SNARE complexes insert GluA1(+) AMPA receptors into the postsynaptic membrane adjacent to spines in an activity dependent manner. Receptors laterally diffuse into the synapse, leading to potentiation B-C) CA1 Stx3 is necessary for novel environment preference B) Novel environment preference assay. During training trials, animals can explore one arm of a T-maze (familiar arm “F”), while the other arm is blocked (novel arm, “N”). During the test trial, the block is removed and animals are free to explore the full maze. ITI - inter-trial-interval. C) CA1 Stx3 is necessary for novel environment preference. *Top* - Representative test trial from a single Cre-injected and a single ΔCre mouse. Heatmaps indicate relative occupancy of each location. *Bottom* - Novel arm preference score [difference score = (time spent in the novel arm) - (time spent in the familiar arm)]. Dotted line indicates equal time spent in the two arms [N=19 control mice, 24 Cre mice. Unpaired t-test: t=2.83, p=0.007] D) Schematic of the head-fixed virtual reality (VR) and two photon (2P) apparatus. E) Task design of the VR novel arm Y-Maze. *Left* - Track schematic. Arrows indicate the animals’ trajectory on left or right trials. Shaded regions indicate reward zones. *Right* - Training protocol. Days 1-5: 5 blocks of “familiar arm” trials (20 trials or 5 minutes, black), 1 block of randomly interleaved “familiar” and “novel arm” trials (40 trials or 10 minutes, orange). Animals are forced to take left or right trajectories on each trial. The animals cannot control the yaw angle in the virtual environment. Day 6: familiar and novel arms randomly interleaved in all blocks. F-G) CA1 Stx3 is necessary for reward anticipation behaviors. F) Control animals (gray) display more anticipatory licking than Cre-injected animals (red). *Left* - peri-reward lick rate on familiar arm trials as a function of position relative to the reward zone on day 5 (magenta - reward zone, shaded regions indicate across animal mean ± sem). *Right* - Average familiar trial peri-reward lick rate on each day. Dots indicate the across trial average for each mouse. Shaded bars indicate across animal mean. [N=9 control mice, 8 Cre mice, 6 days. Mixed effects ANOVA: virus main effect F(1,15)=10.66 p=5.22×10^−4^, day main effect F(5,75)=7.55 p=8.61×10^−6^, interaction F(5,75)=.801 p=.55]. G) Same as (F) for novel arm trials. [N=9 control mice, 7 Cre mice, 6 days. Mixed effects ANOVA: virus main effect F(1,15)=8.40 p=.011, day main effect F(5,75)=2.01 p=.087, interaction F(5,75)=1.33 p=.263] See also Figures S1 & S2.

## Results

### Stx3-dependent postsynaptic membrane fusion in CA1 is critical for novel environment preference and reward anticipation

To investigate the role of Stx3-dependent postsynaptic membrane fusion in forming neural representations within the hippocampus, we first identified a class of behaviors that depends on this molecular pathway. Notably, we did not anticipate that animals lacking CA1 Stx3 would have broad spatial navigation deficits. Mice with a global deletion of GluA1 (i.e. GluA1 knockout animals) do not show gross deficits in a variety of spatial learning tasks^53–55^, and previous work demonstrated that Stx3 is not required in excitatory neurons for spatial learning in a Morris water maze or for contextual fear conditioning^18^. Instead, we hypothesized that CA1 Stx3 may be important for novelty processing. GluA1 knockout animals have specific spatial navigation deficits surrounding short-term memory and novelty processing^54,55^, and decades of theoretical work also posits that CA1 and plasticity in this region play an important role in identifying novelty^56–58^. In addition, a recent study demonstrated that mice with a global mutation impairing GluA1 ubiquitination are deficient in similar assays, suggesting short term memory and novelty processing require regulated AMPAR trafficking^59^. We, therefore, tested whether CA1 Stx3 is required for novel environment preference. We deleted Stx3 in dorsal CA1 neurons by bilateral injection of AAV-Cre in *Stx3*^fl/fl^ mice (“Cre mice”, N=24), with an inactive, mutant version of Cre recombinase serving as a control (AAV-ΔCre, “control mice”, N=19). We achieved nearly complete coverage of the dorsal CA1 region (Fig S1A-B). To probe novelty-guided behavior, we performed a novel-arm T-maze task. Mice were allowed to explore 2 of 3 arms of a T-Maze during training, and the full T-Maze during the test trial (Fig 1B). During the test trial, control mice preferred to explore the previously unseen arm, while Cre-injected mice explored the different arms at chance levels (Fig 1C & S1C). This difference in novelty preference or short-term memory did not appear to be driven by differences in locomotion or anxiety (Fig. S1D-F).

With this behavior in hand, we turned to *in vivo* two-photon Ca^2+^-imaging during head-fixed navigation in a virtual reality (VR) maze. This method enabled us to analyze the role of Stx3 in hippocampal computations by recording the activity of large populations of CA1 neurons as animals performed spatial learning tasks (Fig. 1D). We injected *Stx3*^fl/fl^ mice with a cocktail of viruses in bilateral CA1. All injections contained a virus encoding the Ca^2+^-indicator GCaMP (AAV-hSyn-jGCaMP7f). In one group of animals, the same injection volume included a virus encoding Cre-recombinase and mCherry (AAV-CaMKIIa-mCherry-IRES-Cre, “Cre mice”, N=7, Cre-1 to Cre-7). In the second group of animals, the cocktail included a virus encoding mCherry alone (AAV-hSyn-mCherry, “control mice”, N=9, Ctrl-1 to Ctrl-9). To allow *in vivo* imaging of these neurons, we placed an imaging window over the left dorsal CA1 region. As a conservative measure of Cre recombination, we only analyzed the Ca^2+^-activity of mCherry(+) cells (Fig. S1G-H).

In order to obtain robust recordings of a large population of CA1 neurons while engaging behavioral processes analogous to the freely-moving novel-arm T-maze behavior above, we trained head-fixed mice to perform a VR novel-arm Y-maze task (Fig. 1E & S2A, [Movie S1](https://www.dropbox.com/scl/fi/hyru8d90p7rfoqwbhh3uo/Supplementary-Video1-1080p.mp4?rlkey=q14fq3tbk0m5komaflodhweno&dl=0)). In the freely-moving T-maze task (Fig. 1B-C), Cre and control mice differed significantly in their occupancy of the arms during the test phase. This difference in novel arm preference would complicate the comparison of neural activity across groups. To prevent this potential confound, we controlled the occupancy of the different arms of the virtual Y-maze by forcing mice to take either a left or right “turn” on each trial as they ran forward on a fixed axis treadmill. During training, we forced exploration of 1 of the 2 virtual arms (“familiar”) for 5 blocks of trials while the view of the other arm was blocked with a visual VR curtain. To assess the effects of novel experience on neural coding, while ensuring sufficient exploration of both the familiar and novel arms during the test block, we randomly interleaved trials in which the mouse was forced to run down either the “familiar” or the “novel” virtual arm. We placed unmarked reward zones in distinct locations on each arm (Fig. 1E & S2A, shaded regions) to detect whether mice could distinguish the two arms of the maze. As mice learned the locations of the rewards, they showed distinct reward approach behaviors (e.g. slowing and licking at different locations) on the two arms (Fig. S2B-E). This protocol was repeated for 5 days, and on day 6, mice ran interleaved trials on both arms for the whole session during both training and testing.

While the experimental subjects were not able to express a preference for novelty by choosing which arm to explore, both groups of animals showed behavioral evidence that they recognized novel experiences in this task. In line with our freely-moving novel arm T-Maze, differences in running behavior between the control and Cre mice in block 6 indicate that control animals had a stronger preference for investigating novel stimuli (Fig. S2F-G). In addition, Cre and control mice differed in behaviors related to reward anticipation in this task. Both Cre and control mice rapidly learned the context-dependent reward locations, evidenced by increased licking and decreased running speed only in the spatial bins immediately preceding the reward zones (Fig. 1F-G & S2B-E). Despite evidence of learning in both groups, control mice appeared to have a more reliable reward memory. Control mice licked at a higher rate as they approached the reward zone for both familiar and novel arms (Fig. 1F-G) and approached reward locations with more stereotyped running trajectories (Fig. S2H-I). This difference suggests that abolishing Stx3-dependent membrane fusion may weaken predictive reward coding while sparing the general ability to navigate towards remembered locations.

Novelty and reward are thought to elicit dorsal CA1 plasticity. Intriguingly, prior work showed that novelty and reward correlate with a higher rates of BTSP and SWRs, as well as increased dopamine release^60–70^. To investigate the role of Stx3 in supporting hippocampal computations and plasticity underlying these novelty and reward behaviors, we next turned to the calcium imaging data from this task.

### CA1 Stx3 deletion does not affect gross measures of place code quality and stability

In line with previous reports that animal’s lacking Stx3 do not have general spatial navigation deficits^18^, we observed accurate, reliable, and stable neural representations of space and context in both control and Cre animals. At the level of single trials, we observed repeatable sequences of place cell activity in both control and Cre animals (Fig. 2A-B). Naive Bayesian decoders were highly accurate in decoding the animal’s position from neural activity in both familiar and novel trials, and the decoders’ performance improved as the animals gained experience in the environments over days (Fig. 2A-D). When controlling for the number of cells used in the decoders, we did not find significant differences in decoder error between models trained on data from control or Cre animals (Fig. 2C-D). There was also no difference between groups in the fraction of cells with significant spatial information or the stability of spatial representations within a day (Fig. S3A-B). Place cell codes were largely uncorrelated between the novel and familiar arms of the Y-maze in both groups (i.e. “global remapping”), indicating that contextual discrimination is intact after *Stx3* deletion (Fig. 3E-F & S3C). As a stronger test of intact contextual coding, Cre mice showed normal experience dependent contextual remapping patterns in a separate VR task^71^ (Fig. S4).

**Figure 2.**
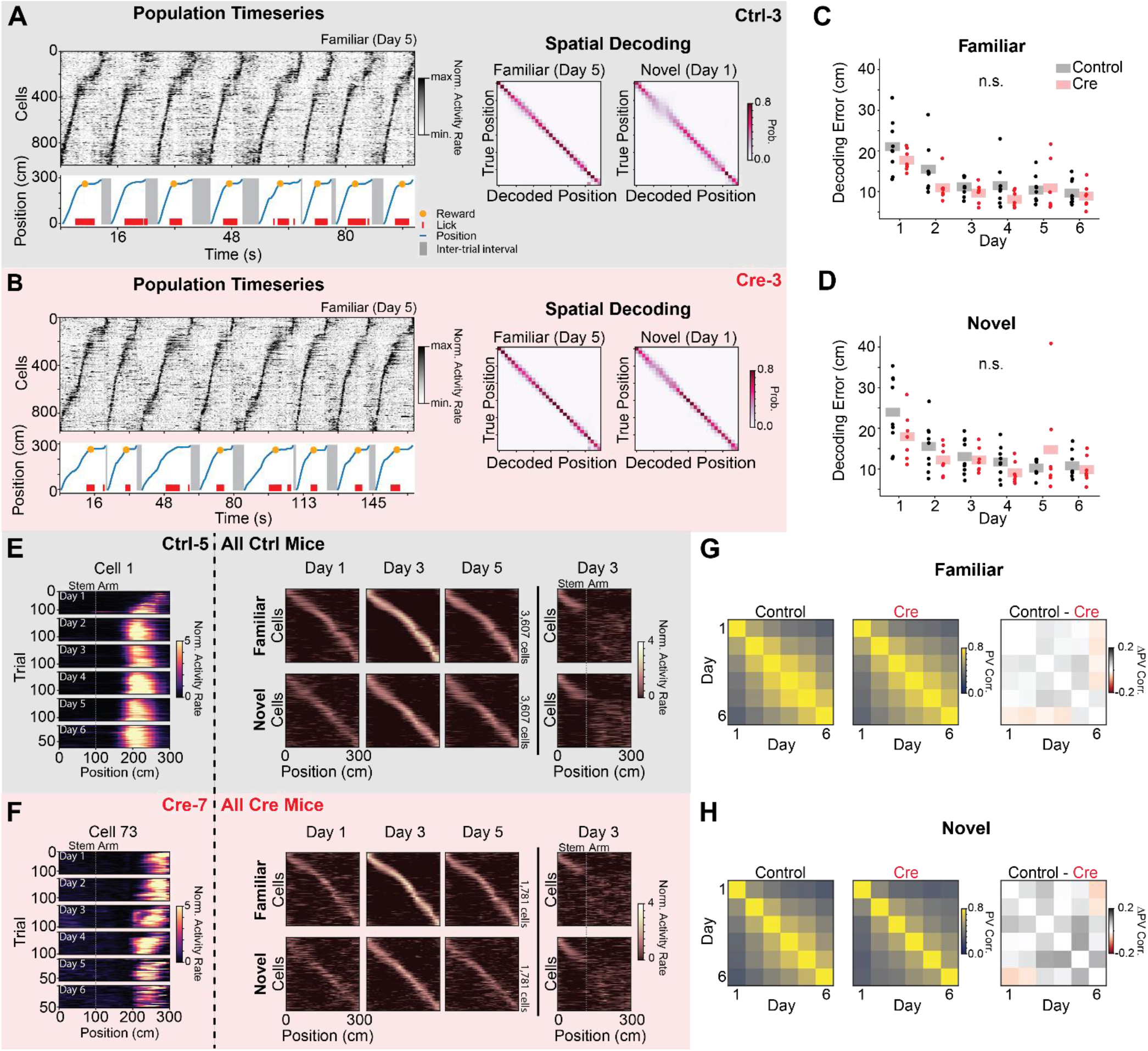
CA1 *Stx3* deletion does not affect gross measures of place code quality and stability. A-D) Accurate neural coding of position does not require CA1 Stx3 A) Example place cell timeseries (“Population Timeseries”*, Left*) and spatial decoding (“Spatial Decoding”, *Right*) from a single control animal (Ctrl-3; gray). *Population Timeseries: Top*-Activity rate of co-recorded place cells over time from a subset of familiar arm trials on day 5. Cells are sorted by their location of peak activity in the average familiar trial activity rate map. *Bottom* - The position of the animal on the virtual track over time (blue). Shaded regions indicate inter-trial intervals. Red dashes are times of licks, and the orange dot indicates when a reward was dispensed. Rewards were randomly omitted on 10% of trials. Note the tight correspondence between the position of the animal on the track and the sequence of neural activity. *Spatial Decoding: Left* - Posterior distributions from leave-one-trial-out naive Bayes decoding of position from familiar trials on day 5 from the same example animal (Ctrl-3, N=1208 cells, 120 trials). *Right* - Same as (*Left*) for novel trials on day 1 (N=1568 cells, 20 trials). Note accurate decoding even on the initial novel trials. B) Same as (A) for a single Cre animal (Cre-3; red). Note similar place cell sequences and decoding accuracy between (A) and (B). (Day 5 familiar trial decoding: N=1087 cells, 17 trials, Day 1 novel trial decoding: N=936 cells, 17 trials) C) Average leave-one-trial out naive Bayes spatial decoding error on familiar trials on each day. Dots indicate the average error of 50 models trained with a random subset of neurons chosen with replacement (n=128 cells per model) for each mouse. Shaded bars indicate across animal mean. [N=9 control mice, 8 Cre mice, 6 days. Mixed effects ANOVA: virus main effect F(1,14)=1.75 p=0.207, day main effect F(5,70)=30.2 p=3.25×10^−16^, interaction F(5,70)=1.70 p=0.145; Posthoc paired t-test (Holm-corrected p-value) comparing decoding error on each day: day 1 vs day 2 - t=8.17 p=7.96 x 10^−6^; day 1 vs day 3 - t=9.26 p=1.76×10-6; day 1 vs day 4 - t=10.2 p=5.79×10^−7^; day 1 vs day 5 - t=6.39 p=1.33×10^−4^; day 1 vs day 6 - t=9.68 p=1.07×10^−6^; day 2 vs day 3 - t=3.35 p=0.039; day 2 vs day 6 - t=3.64 p=.024; all other comparisons p>.05] D) Same as (C) for novel trials. [N=9 control mice, 8 Cre mice, 6 days. Mixed effects ANOVA: virus main effect F(1,14)=.576 p=0.460, day main effect F(5,70)=14.6 p=8.74×10^−10^, interaction F(5,70)=2.96 p=0.017; Posthoc paired t-test (Holm-corrected p-value) comparing decoding error on each day: day 1 vs day 2-control t=5.60 p=6.61×10-4; day 1 vs day 3 - t=5.37 p=9.28×10^−4^; day 1 vs day 4 - t=8.64 p=5.00×10^−6^; day 1 vs day 5 - t=3.38 p=.037; day 1 vs day 6 - t=6.52 p=1.35×10^−4^; day 2 vs day 4 - t=3.67 p=.023; day 2 vs day 6 - t=3.75 p=.021; all other comparisons p>.05 E-H) Comparable stability of spatial coding between Control and Cre animals. E) Activity of place cells tracked over all 6 days from Control animals. *Left* - Position-binned activity rate of a single place cell in familiar trials from all days (mouse: Ctrl-5, cell 1). *Right* - Position-binned trial-averaged activity rate (z-scored) of all place cells tracked over days 1-5 (top row - familiar trials, bottom row - novel trials). Cells are plotted in the same order for each plot within a row and are sorted by their location of peak activity on odd-numbered trials from day 3. Day 3 heatmaps indicate average activity on even-numbered trials. All other heatmaps indicate the average of all trials. These heatmaps show a stable tiling of place cells across days. *Far right* - (top) novel trial activity with cells sorted by familiar trial activity. (bottom) Familiar trial activity with cells with cells sorted by novel trial activity. Place cells that code for the stem of the Y-Maze are visible at the beginning of the track in both conditions. Place cells remap on the arms of the maze as indicated by the disorganized activity rate maps. F) Same as (E) for place cells tracked over all experimental days in Cre animals. G) Across animal average day x day population vector correlation (PV corr.) for familiar trials (*Left:* Control animals, *Middle: Cre* animals, *Right:* difference between groups). Each animal’s population vector is calculated using the trial-averaged spatial activity rate map for each cell on each day. [N=9 control mice, 7 Cre mice. *Right:* control vs Cre unpaired t-test of average across day PV correlation t=.333 p=.745] H) Same as (G) for novel trials. [N=9 control mice, 7 Cre mice. *Right:* control vs Cre unpaired t-test of average across day PV correlation t=1.27 p=.223] See also Figures S3 & S4.

**Figure 3.**
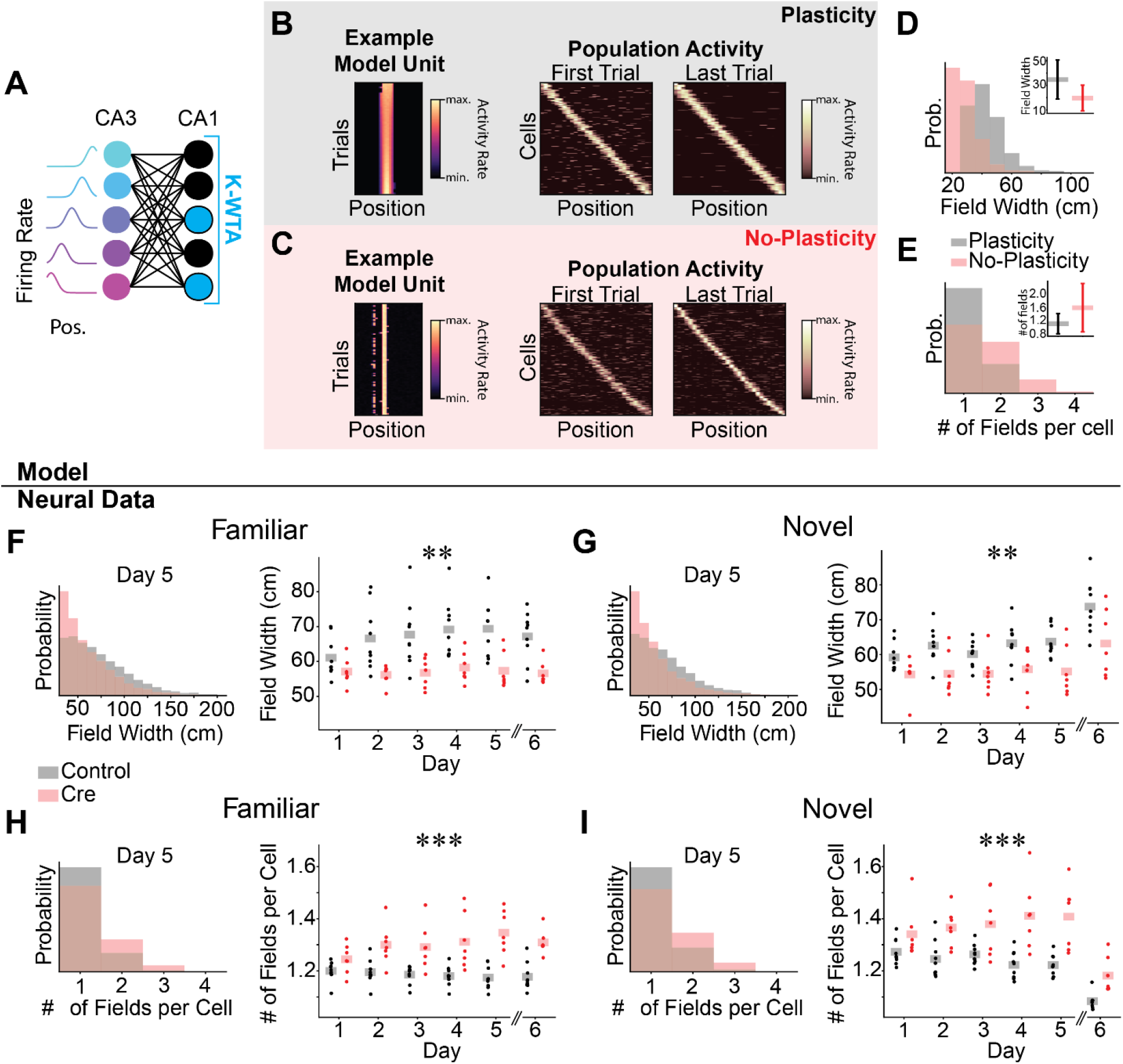
Spatial coding in Cre animals may be passively inherited from upstream activity. A-E) Computational model of the CA3-CA1 circuit. A) Schematic for K-Winners-Take-All (KWTA) model of place field inheritance. A set of place selective cells (“CA3”) projects randomly to a set of output neurons (“CA1”). The set of active neurons at the output layer at each position is determined by a KWTA threshold (the K most excited neurons are active [blue], all other neurons are silent [black]). In the “Plasticity” model, synaptic weights are updated according to a noisy Hebbian rule. In the “No-Plasticity” model, synaptic weights randomly fluctuate. B-C) Both the Plasticity and No-Plasticity models produce stable place cells at the CA1 layer. B) Plasticity model spatial coding. *Left* - Position-binned activity of an example place selective unit from the model. Each row is a single training iteration of the model, akin to a trial in the real experiment. Columns indicate virtual position. *Middle & Right* - The activity of all place selective units is plotted for the first (*Middle*) and last trial (*Right*). Model cells are sorted by their location of peak activity on the last trial. Each row indicates the z-scored activity rate of a unit as a function of position. C) Same as (B) for the No-Plasticity model. Note similar spatial coding and stability in the two models. D) Place fields are wider in the Plasticity model. Histogram of place field widths (black - Plasticity, red - No-Plasticity). *Inset* - mean ± std field width from each model. E) The Plasticity model has fewer place fields per cell than the No-Plasticity model. Histogram of the number of place fields per cell (black - Plasticity, red - No-Plasticity). *Inset* - mean ± std number of fields per cell from each model. F-I) Differences between Control and Cre animal place field properties mirror differences between Plasticity and No-Plasticity models. F-G) Cre animals have narrower place fields. F) Familiar trials. *Left* - Histogram of place field widths for combined data from day 5 (black - Control, red - Cre). *Right* - Within animal average field width on each day. Dots indicate the across cell average for each mouse. Shaded bars indicate across animal means. [N=9 control mice, 7 Cre mice, 5 days (day 6 excluded). Mixed effects ANOVA: virus main effect F(1,14)=10.6 p=.006, day main effect F(4, 56)=4.57 p=0.003, interaction F(4,56)=2.81 p=0.034; day 6 unpaired t-test control vs Cre: t=-3.90 p=.002] G) Same as (F) for novel trials. [N=9 control mice, 7 Cre mice, 5 days (day 6 excluded). Mixed effects ANOVA: virus main effect F(1,14)=10.2 p=.007, day main effect F(4,56)=2.11 p=.091, interaction F(4,56)=.758 p=.557; day 6 unpaired t-test control vs Cre: t=-2.38 p=.036] H-I) Cre animals have more place fields per cell. H) Familiar trials. *Left* - Histogram of number of place fields per cell for day 5 data combined across all mice (Control - black, Cre-red). *Right* - Within animal average number of place fields per cell on each day. Dots indicate the across cell average for each mouse. Shaded bars indicate across animal means. [N=9 control mice, 7 Cre mice, 5 days (day 6 excluded). Mixed effects ANOVA: virus main effect F(1,14)=25.5 p=1.76×10^−4^, day main effect F(4,56)=.804 p=.528, interaction F(4,56)=3.29 p=.017; day 6 unpaired t-test control vs Cre: t=5.30 p=1.19×10^−4^]. I) Same as (H) for novel trials [N=9 control mice, 7 Cre mice, 5 days (day 6 excluded). Mixed effects ANOVA: virus main effect F(1,14)=17.5 p=9.12×10^−4^, day main effect F(4,56)=.158 p=.959, interaction F(4,56)=2.96 p=0.027; day 6 unpaired t-test control vs Cre: t=3.52 p=.008] See also Figure S5.

In order to compare the long-term stability of spatial coding, we identified neurons for which we could reliably track their activity over all six days of the experiment. We observed no obvious differences in the overall stability of place codes between control and Cre animals. Both groups of animals had place cells that maintained constant place coding across each day (Fig. 2E-F), and the population vector correlation of average place cell responses across days did not significantly differ between groups (Fig. 2G-H). Spatial decoders trained across days also did not significantly differ in their accuracy between control and Cre animals (Fig. S3D-G).

While several *in vitro* LTP protocols are intact in animals with reduced Stx3 activity in CA1^17,18^ (but see refs ^14,15^), our results raise the possibility that under certain conditions stable population place codes may not require substantial CA1 synaptic potentiation.

### Spatial coding in Cre animals may be passively inherited from upstream activity

As a proof of concept that place coding in CA1 could be inherited from upstream CA3 population activity without activity-dependent potentiation of CA3-CA1 synapses, we built a simple computational model of the CA3-CA1 circuit (Fig. 3A). In this model, a set of CA3 place cells makes excitatory connections onto a set of CA1 neurons with random weights. To model inhibition in the CA1 layer, a K-Winners-Take-All (KWTA) threshold is applied to CA1 activity. In one model, we allowed CA3-CA1 synaptic weights to update according to a noisy Hebbian rule (“Plasticity” model), and in another model, weights only made small random fluctuations (“No-Plasticity” model). In both models, individual units were clearly place-selective, and the population maintained spatially stable place coding from the first trial (Fig. 3B-C). This model predicts that place representations can emerge in a sparsely active CA1 neural cell population without synaptic potentiation by simply reading out upstream spatially selective inputs. Partially validating a key assumption of our computational model, place fields were intact in the CA3 region and the dentate gyrus in a small cohort of Cre mice in which we additionally injected the same GCaMP virus in these regions (Fig. S5).

Despite stable place coding in both models, we observed several differences in place cell properties between the Plasticity and No-Plasticity models. First, place fields were wider in the Plasticity model than the No-Plasticity model (Fig. 3D). Second, place cells from the No-Plasticity model were more likely to have multiple place fields per cell (Fig. 3E). We observed both of these place coding differences in our *in vivo* calcium imaging data. Place fields from Cre mice were narrower than those from control mice (Fig. 3F-G), and place cells from Cre mice had more place fields on average than place cells from control mice (Fig. 3H-I). Note that these two CA1 place coding properties, narrower place fields and more place fields per cell, have opposing effects on spatial decoder accuracy, perhaps accounting for the comparable decoding performance between control and Cre mice seen in Fig. 2.

Our computational model led us to consider a broader hypothesis. Stx3-dependent place code plasticity in CA1 may not be important for recomputing the place code present upstream in CA3, but perhaps plasticity resulting from Stx3-mediated membrane fusion endows CA1 neurons with additional place coding properties that are unique to CA1 and not present in CA3. Explicitly, we predicted that CA1 neurons from Cre animals should appear more CA3-like in their coding properties. In the following three sections we consider the effect of Stx3 deletion on coding properties that were previously shown to differ between the CA1 and CA3 place cell populations.

### CA1 Stx3 is necessary for increased neural activity in novel environments

The first difference between CA1 and CA3 coding is that CA1 neurons, but not CA3 neurons, increase their firing rates in novel environments^72,73^. Correspondingly, we observed that CA1 neurons in control animals increased their population firing rate in novel environments to a greater extent than Cre animals (Fig. 4A-E & S6). To quantify this effect, we examined the last (6th) block of trials on imaging days 1 through 5. Novel arm trials first appear in the 6th block and are randomly interleaved with familiar arm trials. During this block of trials, we observed an increase in population activity for both familiar and novel arm trials. This increase in activity rate was larger on the arms of the Y-Maze than the stem and greater for novel compared to familiar arm trials in both control and Cre animals. However, control animals showed an overall larger increase in population activity rate than Cre animals in both trial types. The difference in the population activity rate between novel arm and familiar arm trials was also greater in control animals compared to Cre animals. These results were not driven by a general increase in activity rate over the course of imaging or increases in running speed during the last block of trials (Fig. S6 & S2F-G). The magnitude of the increase in activity rate during familiar trials also depended on the time since encountering a novel trial (Fig. S6), suggesting that some of the increase in population activity in familiar trials reflects the interleaved task structure. We also observed a transient increase in population activity at the start of earlier familiar trial blocks on most days, indicating that the start of each block may elicit a small population novelty response (Fig. S6).

**Figure 4.**
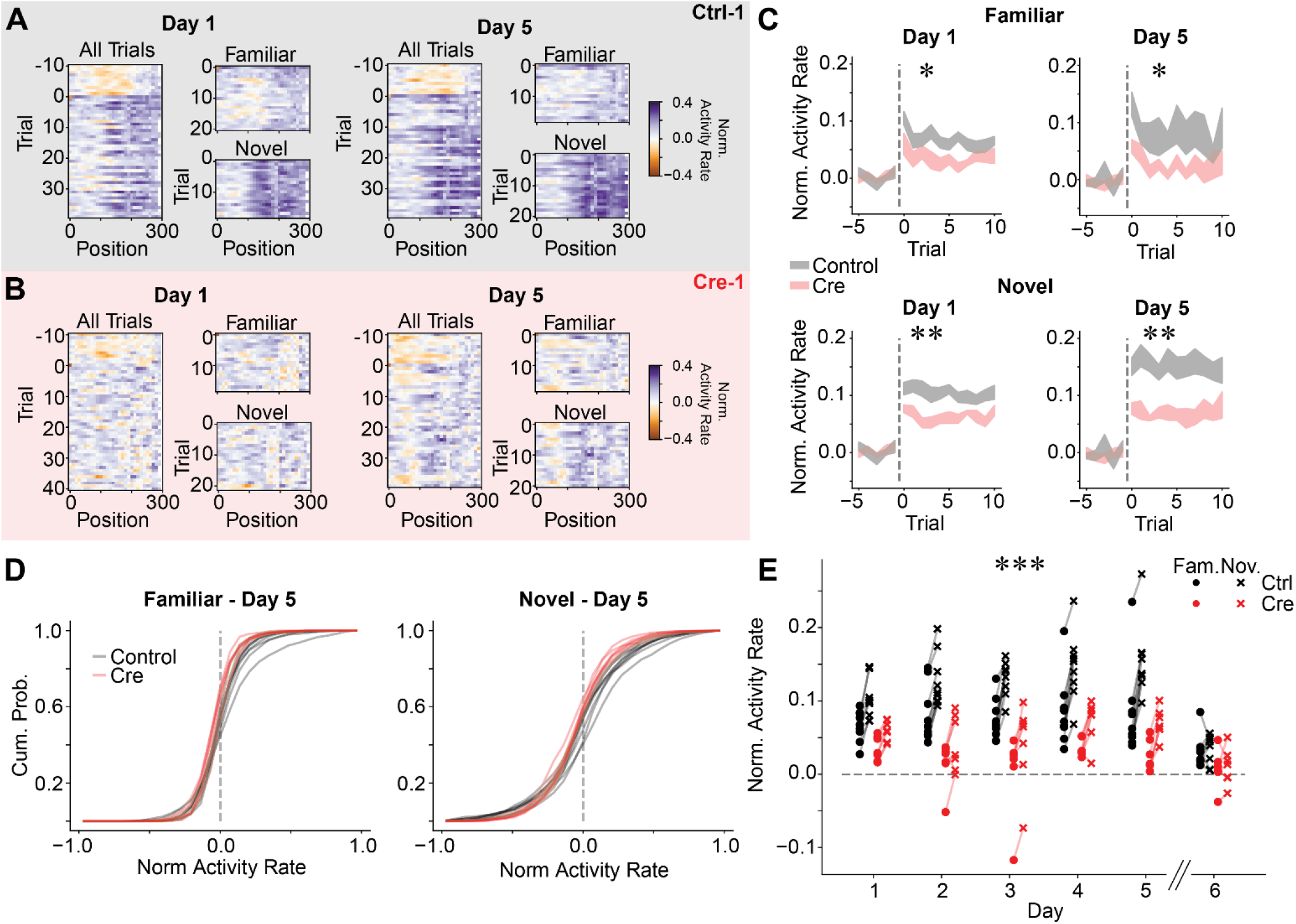
CA1 Stx3 is necessary for novelty-induced increases in neural activity. A) Normalized population activity rate as a function of position on each trial during block 6 for an example control mouse (Ctrl-1). We divide each cell’s activity by its mean activity on the 10 trials preceding the start of block 6 (which occurs at Trial 0). We take the log of the resulting quantity for each cell, so that increases in activity are positive and decreases in activity are negative. This value is plotted as a function of position on each trial. *Left* - Normalized population activity on day 1. Familiar and novel trials are also plotted separately. *Right* - Data from day 5. B) Same as (A) for an example Cre mouse (Cre-1). C) CA1 neural activity increases relative to baseline for both familiar and novel trials during block 6. Cre animals demonstrate a blunted response. Activity rate on each novel trial for days 1 and 5. Data are shown as across animal mean +/− SEM. Dashed line indicates the start of block 6 (*Top-*Familiar trials, *Bottom*-Novel trials). [N=9 control mice, 7 Cre mice, posthoc unpaired t-test of trial-averaged activity rate control vs Cre (Holm-corrected p-value) for mixed effects ANOVA in (E): Familiar Day 1 - t=-3.36 p=.020; Familiar Day 5 - t=-2.73 p=.040; Novel Day 1 - t=-4.04 p=.004; Novel Day 5 - t=-4.69 p=.003] D) Activity rate increases are shared across large portions of the population. Cumulative histograms of trial averaged activity rates across cells on day 5 (*Left* - familiar trials, *Right* - novel trials). Lines indicate histograms from each animal (black - Control, red - Cre). See (C) for information on statistical tests E) Within animal averaged activity rate on block 6 trials (black - Control, red - Cre, dots - familiar trials, “x” - novel trials). Activity rates from Cre animals are significantly lower than activity rates from Control animals for both familiar trials [N=9 Control mice, 7 Cre mice, 5 days (day 6 excluded). Mixed effects ANOVA: virus main effect F(1,14)=16.6 p=.001, day main effect F(4,56)=.944 p=.445, interaction F(4,56)=1.01 p=.412] and novel trials [N=9 Control mice, 7 Cre mice, 5 days (day 6 excluded). Mixed effects ANOVA: virus main effect F(1,14)=37.5 p=2.60×10^−5^, day main effect F(4,56)=4.13 p=.005, interaction F(4,56)=1.60 p=.186]. The difference between novel trial activity and block 6 familiar trial activity is larger in Control animals than Cre animals [N=9 Control mice, 7 Cre mice, 5 days (day 6 excluded). Mixed effects ANOVA: virus main effect F(1,14)=8.13 p=.013, day main effect F(4,56)=6.31 p=2.89×10^−4^, interaction F(4,56)=.443 p=.777]. See also Figure S6.

The increase in activity rate for novel trials did not dissipate over days, and in fact grew stronger over the five days of the experiments (Fig. 4C & E). Examining the cumulative normalized histograms of activity rate in novel trials across cells for each animal, we observed that roughly half of the cells displayed increased activity rates in control animals as compared to Cre animals. Thus, the effect is driven by a large proportion of the population and is not due to a small population of the highly active neurons (Fig 4D). Together these results demonstrate that Stx3 is partially responsible for novelty-induced activity rate increases in CA1.

### CA1 Stx3 is necessary for place cell reward representations and memory

The second difference between CA3 and CA1 coding relates to representations of rewarded locations. Previous works show that a disproportionate number of place cells are recruited to represent goal or reward locations in the dorsal CA1 region but not the CA3 region^74–77^. In CA1, this leads to a population level “overrepresentation” of the spatial locations associated with rewards. In line with this CA3-CA1 difference, we find that control animals allocate a greater proportion of place cells to rewarded locations than Cre animals (Fig. 5A-D & S7). Optogenetically stimulating such reward coding place cells was previously shown to increase licking at non-rewarded locations in a similar VR task^78^, thus the difference in reward representation we observed between control and Cre animals may provide a neural substrate for the matching difference we observed in reward anticipation behaviors (Fig. 1F-G & S2H-I). In addition, control animals but not Cre animals had a significant proportion of “reward cells”, neurons that were active during the approach to reward locations on both arms (Fig. 5A-D)^79^. The proportion of reward cells also increased over days for the control animals indicating accumulating plasticity in the place code (Fig. 5D). This result suggests that Stx3-dependent membrane fusion is important for forming a neural code for salient aspects of the environment.

**Figure 5.**
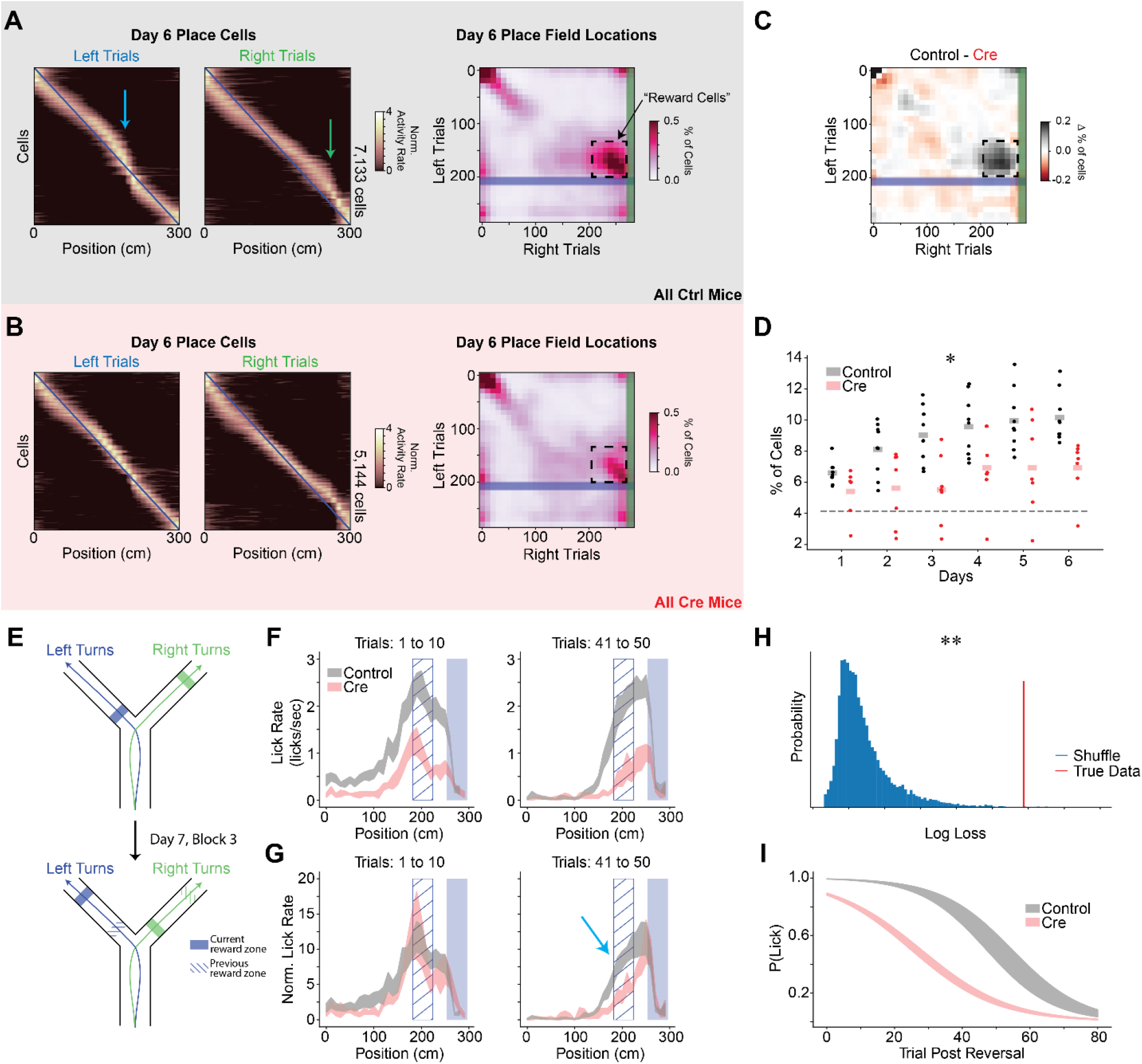
CA1 Stx3 is necessary for reward location memory. A-D) Stx3 is necessary for recruitment of place cells to reward locations. A) *Left* - Day 6 trial-averaged place cell activity (even trials only, odd trial sorted, z-scored) from all control animals. Arrows indicate place cell overrepresentation of rewarded locations. Blue line shows diagonal as a reference. *Right* - Joint normalized histogram of locations of peak activity on left and right trials for place cells on day 6. Rewarded locations are highlighted by the shaded regions (blue - left reward zone, green-right reward zone). Cells that do not remap across trial types will lie on the diagonal. Cells that remap specifically to match shifts in the reward locations will lie on the off-diagonal near the intersection of the reward zones (“reward cells”, dashed box). B) Same as (A) for Cre animals. C) The difference between control and Cre histograms highlights fewer reward cells in the Cre mice. D) Proportion of the population defined as “reward cells” shown for each mouse on each day. Statistical tests performed on logit transformed proportions. [N=9 control, 7 Cre mice, 6 days, Mixed effects ANOVA: virus main effect F(1,14)=8.49 p=.011, day main effect F(5,70)=10.2 p=2.40×10^−7^, interaction F(5,70)=1.46 p=.214]. Dashed line indicates the proportion of reward coding neurons expected by random remapping between novel and familiar arms with constant coding for the stem of the maze. Across-day average fraction of reward cells is greater than chance in control but not Cre animals [One sample t-test: control - t=16.7 p=1.64×10^−7^, Cre - t=2.11 p=.080] E-G) After moving reward locations, Cre animals extinguish licking faster at previously rewarded locations. E) Schematic of the reward reversal task. On day 7, the animals performed the first 2 blocks of trials as in day 6 (randomly interleaved left and right trials). At the beginning of block 3, the reward locations on the two arms were switched. The rewards stayed in this new location for the remaining trials on day 7 and all trials on day 8. F) Lick rate as a function of position for binned left arm trials following the reward reversal (N=9 control mice, 6 Cre mice). Data are shown as across animal mean ± SEM. Solid blue shaded region indicates the current reward zone. Hatched blue region indicates the previous reward zone. G) Normalized lick rate as a function of position for binned trials following the reward reversal. We divide each animal’s lick rate by the mean lick rate on the 10 trials preceding the reversal (N=9 control mice, 6 Cre mice). Data are shown as across animal mean ± SEM. Cyan arrow highlights slower licking extinction at the old reward zone for control mice. H-I) A mixed effects binomial regression was used to quantify the probability of licking more than once in the previous reward anticipatory zone during left arm extinction trials. H) Log loss of model fit (red) vs shuffled distribution (blue). Shuffled distribution was generated by randomly swapping mouse labels of control vs Cre (p=.002). I) Model predictions. Probability of licking more than once in the previous reward anticipatory zone is plotted as a function of trial (black-control, red-Cre). Shaded region shows the best fit model parameters ± estimated standard deviation of the parameters. See also Figure S7.

To test whether the Stx3-dependent overrepresentation of rewards is important for reward-seeking behavior, we switched the reward locations on the arms of the Y-Maze on days 7 and 8 (Fig. 5E). Both groups of animals quickly learned to lick at the new reward locations, but control mice took more trials than Cre mice to extinguish licking at the original reward zone (Fig. 5F-G). We quantified the rate of licking extinction by fitting a binomial generalized linear mixed effects model to the animal’s licking behavior in the old reward zone on left trials. This model estimates the probability that the animal licked more than one time in the previous anticipatory reward zone based on the trial number and whether the animal was in the Cre or control condition. The true model fit exceeded those from a shuffled distribution in which animal identities were scrambled, indicating that the Cre and control animals displayed significantly different licking behavior (Fig. 5H-I). By comparing the individual model parameters to those from the shuffled distribution, we observed a group difference in the intercepts of the binomial regression (shuffled p=.007) but not the slope (shuffled p=.751). Therefore, control animals continued licking at the old reward zone longer than Cre animals, but once control animals began extinguishing their licking for the old reward zone, they did so at a similar rate to Cre animals. Taken together with the differences in reward coding and reward approach behaviors, this difference in licking extinction indicates that CA1 Stx3-mediated place code plasticity is critical for robust reward location memory.

### CA1 Stx3 contributes to experience dependent place field shifts

The third difference between CA3 and CA1 population place codes relates to experience dependent place field location shifts. CA1 place cells tend to shift their firing fields backwards (i.e. opposite the direction of travel) in novel linear environments^80^, which could allow for the formation of predictive codes^38,81,82^. The magnitude of this shift is greater in CA1 neurons than CA3 neurons^83^. A particularly striking version of this backward shift occurs during BTSP. In this form of plasticity, a place field rapidly forms at the location of a dendritic plateau potential^38,44^. Both artificial and spontaneous plateau potentials were previously shown to elicit BTSP^38,41,43,84^. Due to a temporally asymmetric plasticity kernel, the place field shifts abruptly backwards on a linear track following the lap in which the plateau potential occurs^38,39,60^. Recent modeling work shows that gradual shifting of place fields in CA1 is also likely due to repeated bouts of BTSP^85^. Such an abrupt backward shift is unlikely to be inherited from CA3 inputs because BTSP-like plasticity in CA3 utilizes a temporally symmetric plasticity kernel^86^. Consistent with the idea that backward shifting of place fields in CA1 requires synaptic plasticity, we observed a greater extent of rapid place field shifting in novel environments for control animals than Cre animals (Fig. 6 & S8).

**Figure 6.**
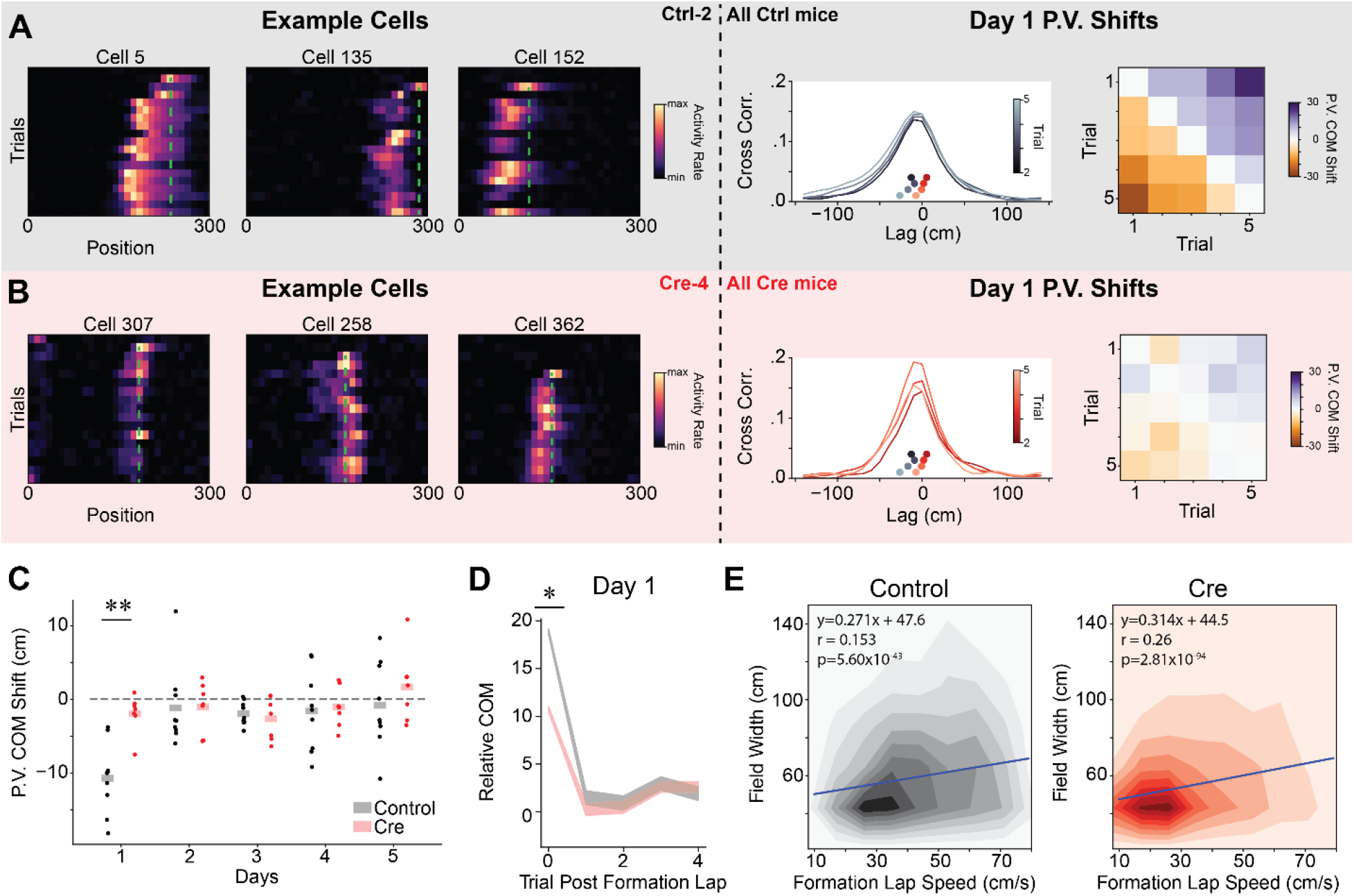
CA1 Stx3 contributes to experience dependent place field shifts in novel environments. A) *Left* - Example co-recorded place cells form a control mouse (Ctrl-2) during novel arm trials on day 1. Cells display an abrupt backwards shift of their place fields after the field first appears. Green dashed line indicates location of peak activity on the place field induction lap. *Middle* - Cross-correlation of population vector activity quantifies population backward shift. For illustration, we show the across animal average population vector (PV) spatial cross-correlation between the first novel trial and the next 4 novel trials on day 1. Inset scatterplot (above x-axis, underneath cross-correlation plot) shows the center of mass (COM) of the cross-correlation (“PV COM shift”) for each trial for both control (greyscale colormap) and Cre (red colormap). Control PV shifts are to the left of the Cre PV shifts indicating a larger magnitude backward shift for control trials. *Right* - Across animal average PV COM shift on day 1 for each pair of initial novel trials. B) Same as (A) for data from Cre animals (example cells Cre-4). Note that smaller abrupt shifts are visible in example cells. C) Average initial novel trial PV shift for each mouse on each day. Dots are the average PV shift for all pairs of initial novel trials on each day for each mouse. Shaded bars are across animal means. [N=9 control mice, 7 Cre mice, 5 days (day 6 excluded). Mixed effects ANOVA: virus main effect F(1,14)=6.41 p=0.024, day main effect F(4,56)=6.09 p=3.85×10^−4^, interaction F(4,56)=2.95 p=0.028. Posthoc unpaired t-test: control vs Cre for day 1-t=-4.63 p=0.002 (Holm-corrected), all other days p>0.05. Posthoc paired t-test (Holm-corrected p-value) for PV shift different from 0: control: day 1 t=-6.66 p=.001, all other days p>.05; Cre: all days p>.05]. D) Individual place fields shift abruptly following their formation lap in novel trials from Day 1. Control animal place fields shift to a greater extent. The COM of place cell activity on each trial, relative to the average place field COM, following the place field’s formation lap. Shaded regions indicate across place field mean +/− SEM (gray-control, red-Cre). After controlling for formation lap running speed through the place field, COM shifts were significantly greater on day 1 for control animals (Linear mixed effects regression, Day 1 - *χ*^2^ =6.37, p=.016). E) The place field width is correlated with running speed through the place field on the formation lap in both groups. Heatmaps show the joint histogram of place field widths and place field formation lap speed through the place field (*Left* - Control, *Right* - Cre). Blue line shows the best fit regression for each condition from mixed effects linear regression. Neither the intercept nor the slope of the regression differs between Cre and control (Linear mixed effects regression, Wald test p>.05) See also Figure S8.

We observed BTSP-like events in CA1 neurons from control animals, particularly on the first day of novel trials (Fig. 6A). As reported previously, control animals displayed widespread population-level backward shifting of place fields in these initial novel arm trials^60,83,85^ (Fig. 6A-C & S8A-C). This effect was not detectable on subsequent days. While we only observed rapid within-day shifting of the place cell code on day 1 novel trials, a slower backward shift of the population place code was also detectable across days during novel arm trials from control animals (Fig. S8G-H). On the other hand, in Cre animals, we observed fewer example cells that show discrete backward place field shifts, and at a population level, there was no evidence of significant population wide backward shifting within a day or across days (Fig. 6B-C & S8). This difference in population markers of rapid plasticity was not driven by group differences in running speed on the initial novel trials (Fig. S8D).

We also quantified rapid place field shifting at the level of single place fields. For each significant place field, we first identified the place field formation lap and then calculated the center of mass (COM) of the cell’s activity within the place field on each subsequent lap. Plotting the COM on each lap, relative to the average place field COM, we observed a discrete backward shift after the formation lap across the population (Fig. 6D & S8E). In agreement with the population vector analysis, the magnitude of the initial place field shift was greater in control animals than Cre animals. This difference remained significant after controlling for running speed through the place field on the formation lap (Fig. S8F). In agreement with previous results^60,85^, we observed a smaller discrete shift in the average place field COM on subsequent days (Fig S8F). This indicates either smaller place field shifts or that BTSP events were less frequent. In line with the BTSP framework, the width of individual place fields correlated with running speed through the place field on the formation lap in both groups of animals (Fig. 6E)^38^. The fact that the correlation of place field width with running speed was preserved but the magnitude of the population COM shift was reduced suggests that Stx3-dependent membrane fusion could be a partial mechanism for BTSP expression.

### CA1 Stx3 is necessary for offline consolidation of place codes

Lastly, we investigated whether Stx3 may contribute to offline consolidation of place codes. Offline consolidation of population codes is thought to largely occur during sharp wave ripples (SWRs), a high frequency local field potential (LFP) oscillation observed in stationary animals during quiet wakefulness and non-REM sleep^46^. During SWRs, sequences of place cells are often “replayed” at highly compressed speeds^45,47,87^. This circuit-level phenomenon may elicit plasticity that stabilizes spatial codes^45,46,51,52,88^.

While we lack LFP data to conclusively identify SWRs, previous work described correlates of SWRs that can be detected using calcium imaging - highly synchronous population activity in CA1 of stationary animals (highly synchronous events, HSEs)^89^. Using a similar approach, we identified HSEs in both control and Cre animals during stationary periods following reward delivery (Fig. 7A & B). There was no difference between groups in the proportion of trials that contained HSEs nor the proportion of cells that participated in each HSE (Fig. S9), indicating that Stx3 is not involved in generating the neural dynamics that initiate and sustain HSEs. However, Stx3 is critical for enhancing the stability of a cell’s spatial coding after it participates in an HSE (Fig. 7A-B). More specifically, in control animals, cells that participated in at least one HSE had higher across day spatial rate map correlation and higher trial x trial rate map correlation than cells that did not participate in HSEs (Fig. 7C-H). Cre animals did not show this boost in spatial coding reliability for cells that participated in HSEs except for a weak increase in across day spatial rate map correlation for novel trials (Fig. 7D-E). The benefit for participating in HSEs was more pronounced in novel environments than familiar environments, consistent with prior work showing that SWRs are more frequent in novel environments^47,62,63^. While there is no difference between control and Cre animals in the population level spatial coding stability (Fig. 2 & S2), this analysis demonstrates that the smaller population of cells that participate in HSEs are selectively stabilized in a Stx3-dependent manner.

**Figure 7.**
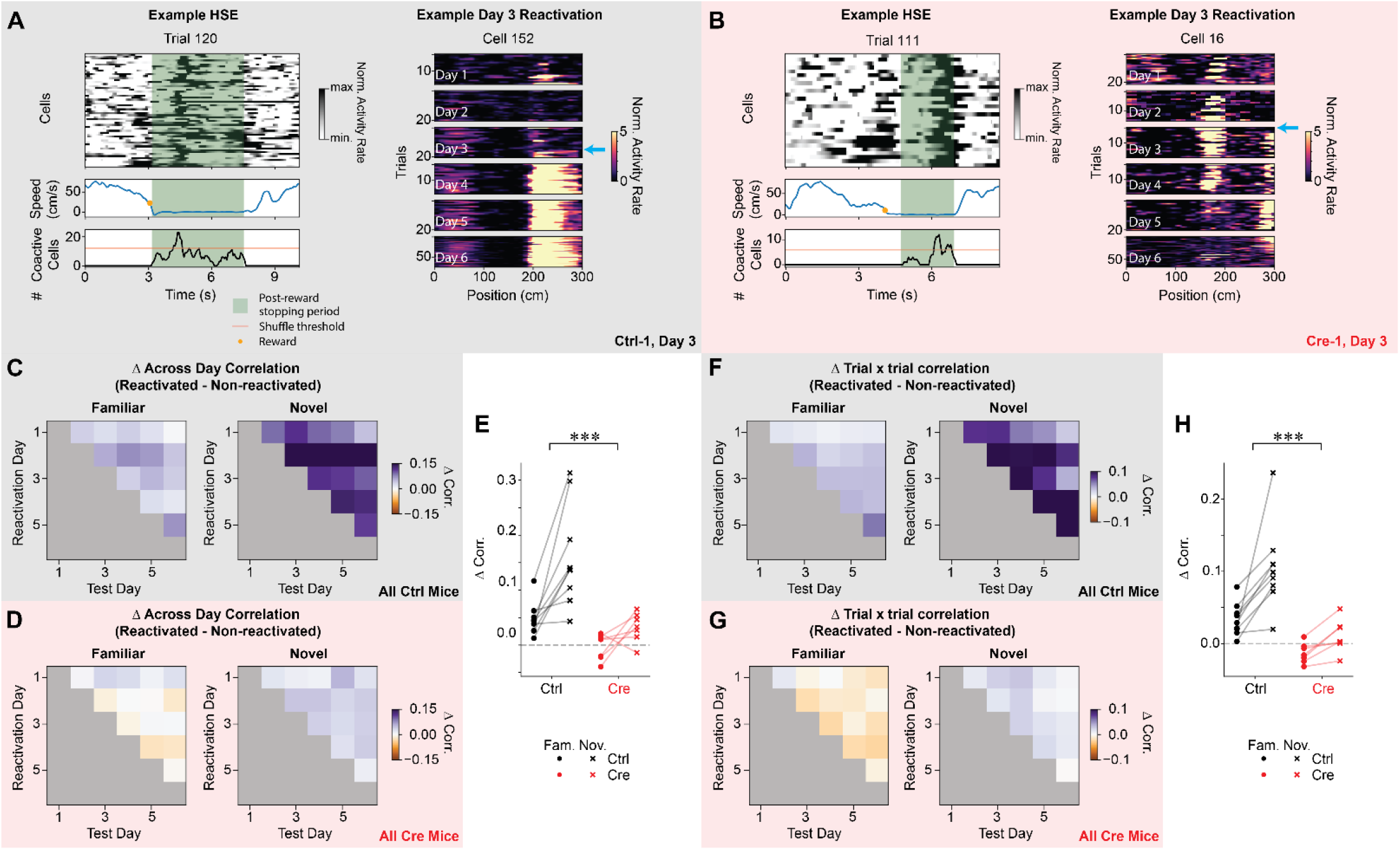
CA1 Stx3 is necessary for offline consolidation of place codes. A) *Left* - Example highly synchronous event (HSE) from a novel trial from a control animal (Ctrl-1) on day 3. *Top* - Activity rate of cells that participate in the HSE over time on a single trial. *Middle* - Running speed. Orange dot indicates the time of reward delivery. *Bottom* - Number of simultaneously active neurons in the population during the post-reward stopping period. Orange line indicates threshold for defining a HSE from shuffled data. Green shading indicates post-reward stopping periods eligible for HSE analysis. *Right* - Example novel arm place cell that is reactivated during a HSE for the first time on day 3 from a control animal (Ctrl-1). The cyan arrow highlights the trial of the first HSE in which the cell participates. Note that the place field appears more stable and reliable on subsequent days. B) Same as (A) for a Cre animal (Cre-1). *Right* - Note that the place cell destabilizes on subsequent days. C-E) For control but not Cre animals, cells that are reactivated during HSEs are more stable across days than cells that are not reactivated during HSEs. This effect is greater in novel trials. C) Difference in across day spatial rate map correlation between reactivated and non-reactivated cells for control animals in familiar and novel trials. Data shown as across animal average. D) Same as (C) for Cre animals. E) Average difference in across day correlation between reactivated and non-reactivated cells. Each point is the average difference in correlation for each mouse (dots - familiar trials, “x” - novel trials, black - control, red - Cre). Reactivated cells in control animals have higher across day correlation than non-reactivated cells for both familiar (one-sample t-test: t=4.67 p=.005, all p-values are Holm-corrected for multiple comparisons) and novel trials (one-sample t-test: t=5.23 p=.003). In Cre animals, reactivated cells have higher across day correlation for novel trials (one-sample t-test: t=3.23 p=.036) but not for familiar trials (one-sample t-test: t=-.367 p=.726). The increase in across day correlation for reactivated cells is greater for control animals than Cre animals. This increase in stability of the neural representation is larger for novel trials than familiar trials [N=9 control mice, 7 Cre mice. Mixed effects ANOVA: virus main effect F(1,14)=18.1 p=7.94×10^−4^, familiar vs novel main effect F(1,14)=20.7 p=4.54×10^−4^, interaction F(1,14)=4.91 p=0.044]. Furthermore, the effect of novelty is greater for control animals [Novelty effect control vs Cre: unpaired t-test t=2.22 p=.044]. F-H) For control but not Cre animals, cells that are reactivated during HSEs have higher trial by trial correlation than cells that are not reactivated. This effect is greater in novel trials. F) Difference in trial-by-trial spatial rate map correlation between reactivated and non-reactivated cells for control animals in familiar and novel trials. Data shown as across animal mean. G) Same as (J) for Cre animals. H) Average difference in trial-by-trial correlation between reactivated and non-reactivated cells. Each point is the average difference in correlation for each mouse (dots - familiar trials, “x”-novel trials, black - control, red - Cre). Reactivated cells in control animals have higher trial x trial correlation than non-reactivated cells for both familiar (one-sample t-test: t=4.46 p=.006, all p-values Holm-corrected for multiple comparisons) and novel trials (one-sample t-test: t=5.44 p=6.12×10^−5^). In Cre animals, reactivated cells do not have higher trial x trial correlation than non-reactivated cells for either trial type (Holm-corrected p>.05). The increase in trial-by-trial correlation for reactivated cells is greater for control animals than Cre animals. This increase in stability of the neural representation is larger for novel trials than familiar trials [N=9 control mice, 7 Cre mice. Mixed effects ANOVA: virus main effect F(1,14)=25.9 p=1.66×10^−4^, familiar vs novel main effect F(1,14)=20.5 p=4.76×10^−4^, interaction F(1,14)=4.47 p=0.053]. See also Figure S9.

## Discussion

In this work we link the postsynaptic membrane fusion machinery implicated in regulated exocytosis of postsynaptic components, principally AMPARs, with CA1 computations that support behavior. In particular, we find that Stx3 is necessary for forming predictive codes of space in novel environments, strengthening spatial reward memories and selectively stabilizing the place codes of neurons that participate in offline consolidation events. We also find that Stx3-mediated membrane fusion plays a role in shaping CA1 place fields, but that this pathway for postsynaptic membrane fusion was not critical for maintaining an overall coherent place code. Together these results point to activity dependent postsynaptic membrane trafficking in CA1 as a mechanism for enriching presynaptic contextual and place codes with salient and predictive information about an animal’s environment, without the need to recompute the information already present in the presynaptic inputs.

### Enriching and stabilizing hippocampal place codes through Stx3-dependent plasticity

In agreement with previous theoretical and experimental evidence, our results suggest that Stx3-dependent synaptic plasticity in dorsal CA1 incorporates and consolidates predictive and salient information into the hippocampal place code, allowing animals to anticipate rewards and identify novel stimuli^40,56,81,90–92^. A number of different plasticity phenomena have been implicated in the processing of rewarded and novel stimuli, suggesting that Stx3-dependent postsynaptic membrane fusion may be a common molecular mechanism for these different forms of plasticity.

Perhaps the simplest form of Stx3-dependent spatial coding we observed was that CA1 neurons from control animals increased their activity in novel environments to a greater extent than Cre animals (Fig. 4). This increase in CA1 population activity in control animals may partially underlie the corresponding preference for novel stimuli we observed in the control animals. Though we do not directly observe GluA1(+) AMPAR dynamics in this work, this result suggests that the CA1 activity rate increase in novel environments^72,73^ is driven by Stx3-dependent insertion of new AMPAR into the postsynaptic membrane. This Stx3-dependent novelty signal may also contribute to the increased rate of BTSP-like place field formation events seen in novel environments for control animals^60,84^. However, it is unlikely that BTSP itself drives the overall activity rate increase because this effect outlasts the period in which most BTSP-like events occur^60^.

By analyzing the backward shifting of place fields in novel environments, we demonstrated that Stx3 is necessary for full expression of *in vivo* correlates of BTSP-like plasticity. The abrupt backward place field shifts in BTSP are a result of a temporally asymmetric learning rule^38^. Meaning that after a BTSP-inducing plateau potential, CA1 neurons become active in anticipation of the CA3 neurons that originally drove them to spike (i.e. they form a predictive code)^40^. It was recently confirmed that BTSP is expressed by changes in synaptic efficacy between CA3 and CA1 neurons *in vivo*^42^. BTSP also requires CaMKIIa signaling, suggesting that canonical LTP expression mechanisms may underlie BTSP^48,49^. Our work extends these findings and demonstrates that full expression of BTSP may require specific membrane fusion proteins implicated in regulated GluA1(+) AMPAR exocytosis. One possible explanation for this result is that rapid potentiation during BTSP-like events depletes the pool of extrasynaptic AMPARs that can readily diffuse into the postsynaptic density before the synapses reach their full desired strength. Stx3-dependent membrane fusion replenishes this pool of AMPARs to allow for complete plasticity expression. Future work will need to investigate the exact role Stx3 plays in BTSP expression.

In addition to their roles in BTSP, NMDARs and entorhinal cortex inputs that drive dendritic plateau potentials are necessary for forming CA1 place cell overrepresentations of rewarded locations^61,75^. Our finding that Stx3 is also necessary for forming representations of rewarded locations provides corroborating evidence that reward representations in CA1 place codes require synaptic plasticity^68,75,93–95^. A parsimonious explanation is that Stx3’s potential role in BTSP expression mediates the necessity of Stx3 in reward overrepresentations. However, rewards are also associated with dopamine release from ventral tegmental area (VTA) and locus coeruleus (LC) axons in the dorsal hippocampus^66–69^. Stimulation of LC axons in the presence of reward is sufficient to drive overrepresentation of rewarded locations, and inhibiting LC axons or VTA neurons prevents recruitment of additional place cells to rewarded locations^68,69^. It is currently unclear if this effect is due to dopamine increasing the rate of BTSP events or if dopamine also stimulates other plasticity mechanisms in which Stx3 may be involved^67,96,97^.

Dopamine release also partially determines which neural representations will be reactivated during another plasticity-associated event, SWRs^98^. Offline reactivation of sequences of place cells during SWRs is necessary for the performance of a variety of memory-guided tasks and for stabilizing hippocampal codes^99–105^. Reactivation during SWRs is particularly important to the stabilization of place codes in novel environments and of place cell representations away from rewarded locations^89,106^. Here, we show that the selective stabilization of cells that participate in HSEs, a population event strongly associated with SWRs, is abolished by *Stx3* deletion. Taken together, our results suggest that activity-dependent membrane trafficking may be a common mechanism for expression of a variety of behaviorally relevant plasticity phenomena including BTSP and SWRs.

### Intact place coding and navigation behavior in animals lacking Stx3

Animals lacking CA1 Stx3 had accurate and spatially stable place codes that downstream brain regions could potentially use to read out the animal’s location in space (Fig. 2 & S3). This result provides potential insight as to why some spatial learning behaviors are intact in Stx3-deficient animals. One possible interpretation is that Stx3 is not necessary for *in vivo* plasticity that supports place code formation. However, our modeling results show that place codes can pass through random CA3-CA1 connectivity without any plasticity. In the model, plasticity in CA1 place cells causes increased place field width and fewer place fields per cell (Fig. 3). The intuition for this result is that random initial biases in synaptic connectivity drive some CA1 neurons to be more excited than others at each location in space. It is unlikely that the same population of CA1 neurons is the most excited at multiple locations, so the K-winners-take-all activity threshold yields a unique population of CA1 neurons activated at each location in space (i.e. a stable CA1 place code). When plasticity is added to the model, the CA3-CA1 synapses that initially drove each CA1 cell to fire are strengthened. This exaggerates the random biases in connectivity and subsequently drives the CA1 cell to be active at adjacent locations within the CA3 neuron’s place field^84^. Consequently, the CA1 neuron’s place field widens. As a result of the widening place fields, the least active CA1 neurons at these expanded locations no longer cross the K-winners-take-all threshold and drop out of the place cell code. This leads to a reduction in the average number of place fields per cell. Consistent with the model, we observed wider place fields and fewer place fields per cell in the control animals than the Cre animals, demonstrating that synaptic potentiation is most likely reduced in the Cre animals. Our results further validate several single cell models predicting that LTP broadens place fields^38,81^. While we emphasize the role of CA3-CA1 plasticity in the context of this study, plastic entorhinal cortex-CA1 synapses at the distal tuft dendrites of CA1 neurons and recurrent CA1-CA1 synapses play important roles in determining the coding properties of CA1 neurons^43,107^.

Our hypothesis that a CA1 place cell can inherit spatial selectivity from upstream inputs without synaptic potentiation is also consistent with previous findings: place cells receive more presynaptic inputs inside their naturally occurring place fields^108^, and increasing the excitability of a cell can unveil “hidden” place fields^109^. Although CA1 pyramidal cells appear equipped with the synaptic inputs and plasticity mechanisms to become a place cell at any location^39,41–43,110^, neural dynamics alone may be sufficient for the CA1 population to read off a place code from its upstream inputs.

Our findings also fit with previous work that shows place coding can exist without typical functioning of other critical components of postsynaptic plasticity machinery such as NMDA receptors, GluA1, or CaMKIIa^110–114^. In all of these mutant backgrounds, spatial coding is present but mutant animals have differences in precision, stability and reliability of place coding, particularly in novel environments. While these important works clearly demonstrate the necessity of proteins involved in LTP for place code plasticity, these manipulations are not specific to LTP *per se*. GluA1, NMDA receptor, and CaMKIIa mutant animals have deficits in synaptic and dendritic integration^53,115–119^, making it ambiguous whether place coding deficits in these animals are due to a lack of synaptic plasticity or an inability of the cell to properly integrate its inputs^116,120,121^. NMDA receptors and CaMKIIa are also implicated in broad plasticity pathways including structural plasticity, modifications to the postsynaptic density, and posttranslational modification of AMPARs^1,122–125^. Deleting CA1 Stx3 is a more subtle and specific manipulation that leaves basal synaptic integration intact and is believed to predominantly affect exocytosis of GluA1(+) AMPARs in response to LTP-inducing stimuli. Stx3(-) membrane fusion complexes are likely capable of delivering many of the same synaptic components to the membrane but without the same temporal control. Consequently, Stx3 deletion is unlikely to completely abolish all forms of synaptic potentiation that rely on postsynaptic membrane fusion.

In agreement with this interpretation, our results largely indicate a partial loss of wildtype CA1 plasticity phenotypes in Cre animals. Cre animals increase CA1 population activity rates in novel environments, but to a lesser extent than control animals (Fig. 4). Cre animals have a larger population of reward coding neurons than expected by chance but fewer than control animals (Fig. S7). Place fields from Cre animals show a significant abrupt backward shift of place field firing in novel environments (Fig. S8), but this shift is smaller than those from control animals and too small to observe at the level of population vector shifts (Fig. 6). Participating in HSEs weakly improved across day spatial coding stability in Cre animals, but to a lesser extent than in control animals (Fig. 7E).

## Conclusion

Activity-dependent membrane trafficking at postsynaptic specializations through dedicated molecular pathways remains a dominant model for synaptic potentiation. There is a wealth of *in vitro* data supporting this hypothesis^1,4^ and an increasing amount of *in vivo* data demonstrates that learning is associated with increased surface expression of GluA1(+) AMPARs^130,131^. However, few experiments have linked the regulation of postsynaptic membrane traffic to neural computations *in vivo*. Here, we took advantage of the well-studied spatial and contextual representations of the hippocampal circuit to demonstrate how activity-dependent postsynaptic membrane trafficking may allow neural circuits to layer computations through sequential steps of processing. We hope that this work can provide a framework for understanding how redundant neural representations found across the brain could act as a substrate for plasticity^132–134^ and a model for how to isolate the unique contribution of synaptic plasticity expression mechanisms within a brain region.

## Acknowledgments

We thank A. Diaz, M. Sosa, D. Luna, G. Wang, and L. Ho for help with animal care and training.

National Institute of Health Grants 1R01MH126904-01A1 (LMG), R01MH130452 (LMG), BRAIN Initiative U19NS118284 (LMG), The Vallee Foundation (LMG), The James S. McDonnell Foundation (LMG), The Simons Foundation 542987SPI (LMG) The Scully Project (TCS), The Howard Hughes Medical Institute (TCS)

This article is subject to HHMI’s Open Access to Publications policy. HHMI lab heads have previously granted a nonexclusive CC BY 4.0 license to the public and a sublicensable license to HHMI in their research articles. Pursuant to those licenses, the author-accepted manuscript of this article can be made freely available under a CC BY 4.0 license immediately upon publication.

## Author Contributions

Conceptualization (MHP, KK, TCS, LMG), Methodology (MHP, KK, TCS, LMG), Investigation (MHP, KK), Visualization (MHP, KK), Funding acquisition (TCS, LMG), Project administration and supervision for freely moving behavioral experiments (TCS), Project administration and supervision for two photon and virtual reality experiments (LMG), Writing - original draft (MHP, KK), Writing - review and editing (MHP, KK, TCS, LMG)

---

**Figure S1. Related to Figure 1.**
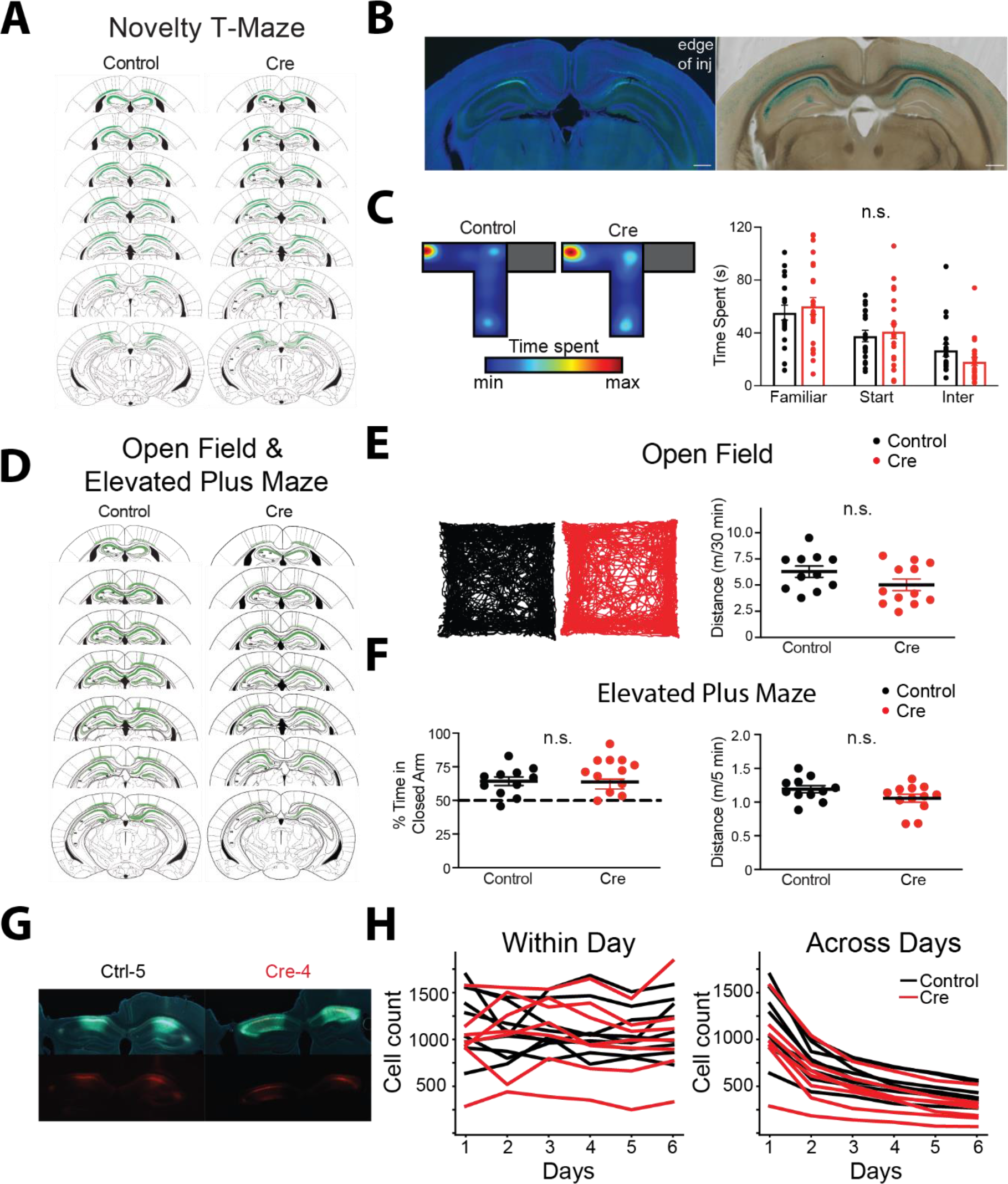
A) Summary plot of viral spread for every mouse that underwent the Novelty T-Maze. Each mouse is drawn with a semi-transparent fill – darker green indicates more overlap across mice. B) Quantification of viral spread based on EGFP fluorescence underestimates extent of Cre recombination. Consecutive 100 µm thick sections of a Cre-injected *Stx3^fl/fl^* mouse at the posterior edge of EGFP expression. *Left* - EGFP (green) with DAPI counterstain (blue), *Right* - Ꞵ-galactosidase chromogenic stain. Ꞵ-galactosidase is only expressed upon Cre recombination. Scale bars: 500 µm C) *Left* - Color-coded occupancy plot of a representative mouse from each group during novelty T-Maze training. *Right* - CA1 deletion of *Stx3* does not affect novelty T-Maze training, as demonstrated by the summary plot of time spent in training T-Maze compartments (averaged over all trials for each mouse). Control and Cre mice occupied the 3 zones of the maze during training at similar rates. Dots represent occupancy for each mouse. Bars indicate across animal mean. “Inter” = intermediate zone, the zone at the intersection of all arms. Dots indicate each mouse. Bars indicate across animal mean. Error bars indicate ± SEM. [N=19 control mice, 24 Cre mice. rmANOVA: compartment main effect F(1.565, 64.18)=15.3, p<1.00-5, virus main effect F(1,41)=.061, p=.806, interaction F(2,82)=.681 p=.509]. D) Same as (A) for every mouse that underwent open field and elevated plus maze training. E) *Left* - Representative open field tracks. *Right* - CA1 deletion of *Stx3* does not affect basal locomotion as demonstrated by the summary plot of distance traveled in the open field. Dots indicate each mouse. Lines indicate across animal mean. Error bars indicate ± SEM. Dots represent traveled for each mouse. Bars indicate across animal mean. Error bars are SEM. [N=11 control mice, 12 Cre mice. Unpaired t-test t=1.64 p=0.116] F) *Left* - CA1 deletion of *Stx3* does not affect anxiety-like behavior as demonstrated by the summary plot of %time spent in the closed arm of the elevated plus maze. Dots indicate each mouse. Lines indicate across animal mean. Error bars indicate ± SEM. Dotted line indicates chance occupancy. [N=11 control mice, 12 Cre mice. Unpaired t-test: t=.096 p=.924] *Right* - CA1 deletion of *Stx3* does not affect locomotion in the elevated plus maze as demonstrated by the summary plot of distance traveled. Dots represent the behavior of each animal. Bars indicate across animal means. Error bars are SEM. [N=11 control mice, 12 Cre mice. Unpaired t-test: t=1.68 p=.107]. G) Histology from two example mice(*Left* - Ctrl-5, *Right* - Cre-4). Coronal slice showing unilateral imaging implant and bilateral virus expression. Top – Co-expression of DAPI (blue), GCaMP (green), and mCherry (red). Bottom - Expression of mCherry. H) *Left* - Number of cells recorded in each session for each mouse (black - control, red - Cre). Only GCaMP^+^/mCherry^+^ neurons were included in all analyses. *Right* - Number of cells tracked across all sessions using ROIs from day 1.

**Figure S2. Related to Figure 1.**
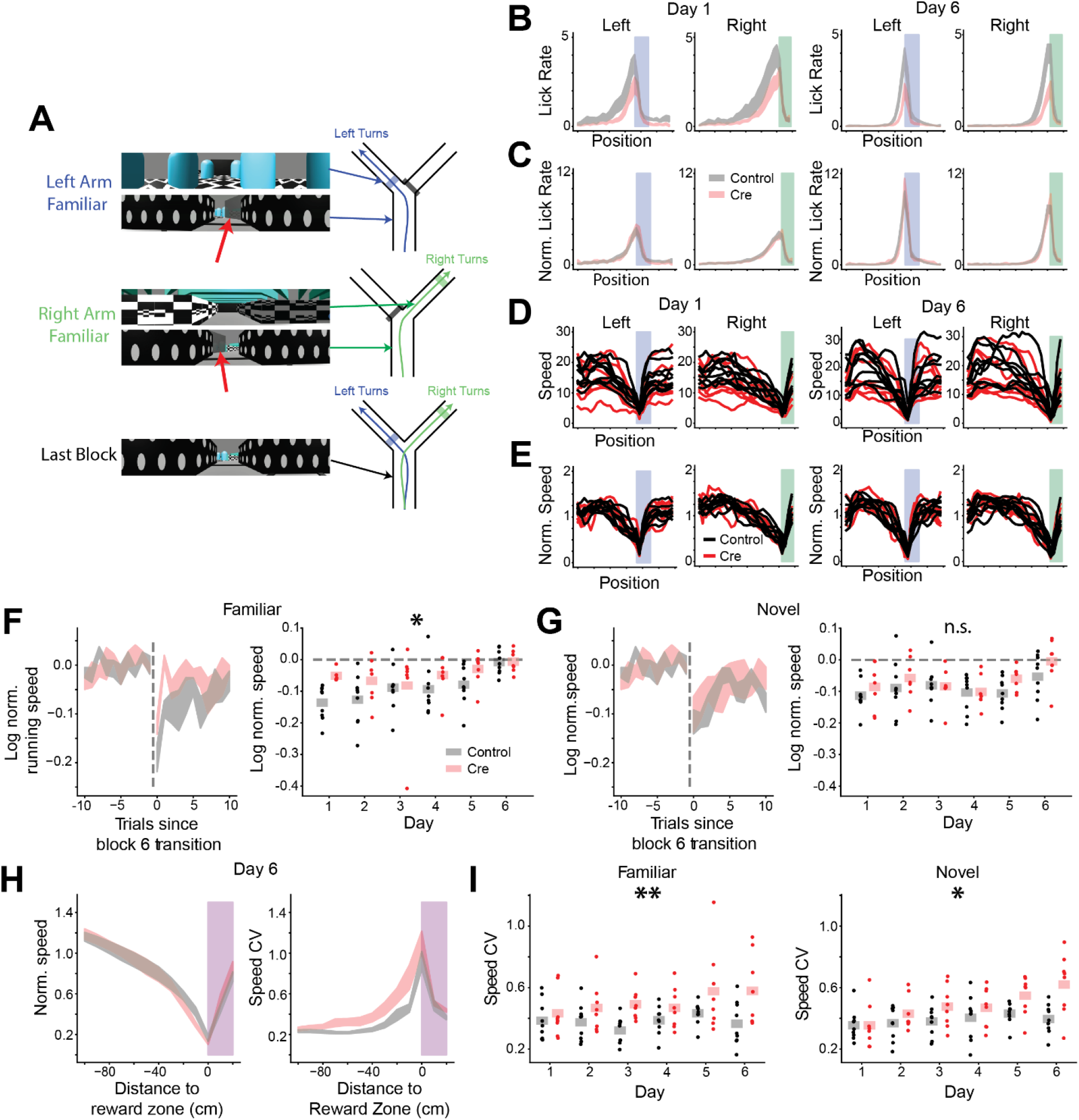
A) Example screenshots of virtual reality track for left arm familiar trials (*Top*), right arm familiar trials (*Middle*) and the stem of trials from the last block where the virtual curtain is removed (*Bottom*). Red arrows highlight the location of the VR curtain during training. We counterbalanced whether the right or left arm was the familiar arm across animals. B) Mean lick rate as a function of position for left and right trials for days 1 and 6. Reward zones are highlighted as in (A) and Fig. 1E. This serves as a control analysis to ensure across group differences in lick rates are not driven by differences across the virtual environments. Novel and familiar trials are intermixed here (black - control, red - Cre, shaded regions indicate across animal mean ± SEM). C) Mean normalized lick rate as a function of position for left and right trials for days 1 and 6. Each mouse’s position-binned lick rate is divided by the mouse’s overall average lick rate. Reward zones are highlighted as in (B). (black - control, red - Cre, shaded regions indicate across animal mean ± SEM. D) Mean running speed as a function of position for left and right trials for days 1 and 6 (black - control, red - Cre, each line is a different mouse). E) Mean normalized running speed as a function of position for left and right trials on days 1 and 6. Each mouse’s position-binned running speed is divided by the mouse’s overall average running speed. The characteristic decrease in running speed to consume the reward can be seen most clearly here. (black - control, red - Cre, each line is a different mouse). F-G) Cre animals react less to visual novelty in VR. F) Control animals slow down more than Cre animals in familiar arm trials after the novel arm block is removed. *Left* - Normalized running speed is plotted for the familiar arm trials surrounding the transition to block 6 on day 1 (red - Cre, black-control, dotted line-transition between blocks 5 and 6). Data shown as across animal mean +/− SEM. *Right* - Average normalized running speed for the first two familiar arm trials of block 6. Dots indicate each mouse. Shaded bars indicate across animal mean. [N=9 control mice, 7 Cre mice, 5 days (day 6 excluded). Mixed effects ANOVA: virus main effect F(1,14)=4.82 p=0.045, day main effect, F(4,56)=1.14 p=.348, interaction F(4,56)=.687 p=.604; posthoc unpaired t-tests for virus effect on each day, day 1 Holm-corrected t=4.43 p=.007, days 2-6 p>.05] G) Control and Cre animals slow similarly to novel arm trajectories. Same as (F) for novel arm normalized running speed. [N=9 control mice, 7 Cre mice, 5 days (day 6 excluded). Mixed effects ANOVA: virus main effect F(1,14)=1.12 p=.308, day main effect F(4,56)=.802 p=.529, interaction F(4,56)=.603 p=.662] H-I) Control animals show more stereotyped reward approach trajectories than Cre-injected animals. H) Control animals have more stereotyped reward approach trajectories despite comparable average running profiles. *Left* - On average, running speed as a function of position is comparable between control and Cre animals. Normalized running speed as a function of position relative to the reward zone for all trials on day 6. *Right* - Cre animals have more variable running speed during reward approach indicated by a higher coefficient of variation (CV, standard deviation / mean) of running speed in the spatial bins preceding rewards. CV of running speed across all trials on day 6 as function of position relative to reward. Shaded regions indicate mean +/− SEM. I) Average peri-reward running speed CV on each day for familiar (*Left*) and novel (*Right*) trials. Dots indicated across trial average for each mouse. Shaded bars indicate across animal mean. [N=9 control, 7 Cre mice, 6 days, mixed effects ANOVA. Familiar trials: virus main effect F(1,15)=9.99 p=.006, day main effect F(5,75)=1.33 p=.262, interaction F(5,75)=.853 p=.517. Novel trials: virus main effect F(1,15)=6.10 p=.026, day main effect F(5,75)=4.41 p=.001, interaction F(5,75)=2.06 p=.080]

**Figure S3. Related to Figure 2.**
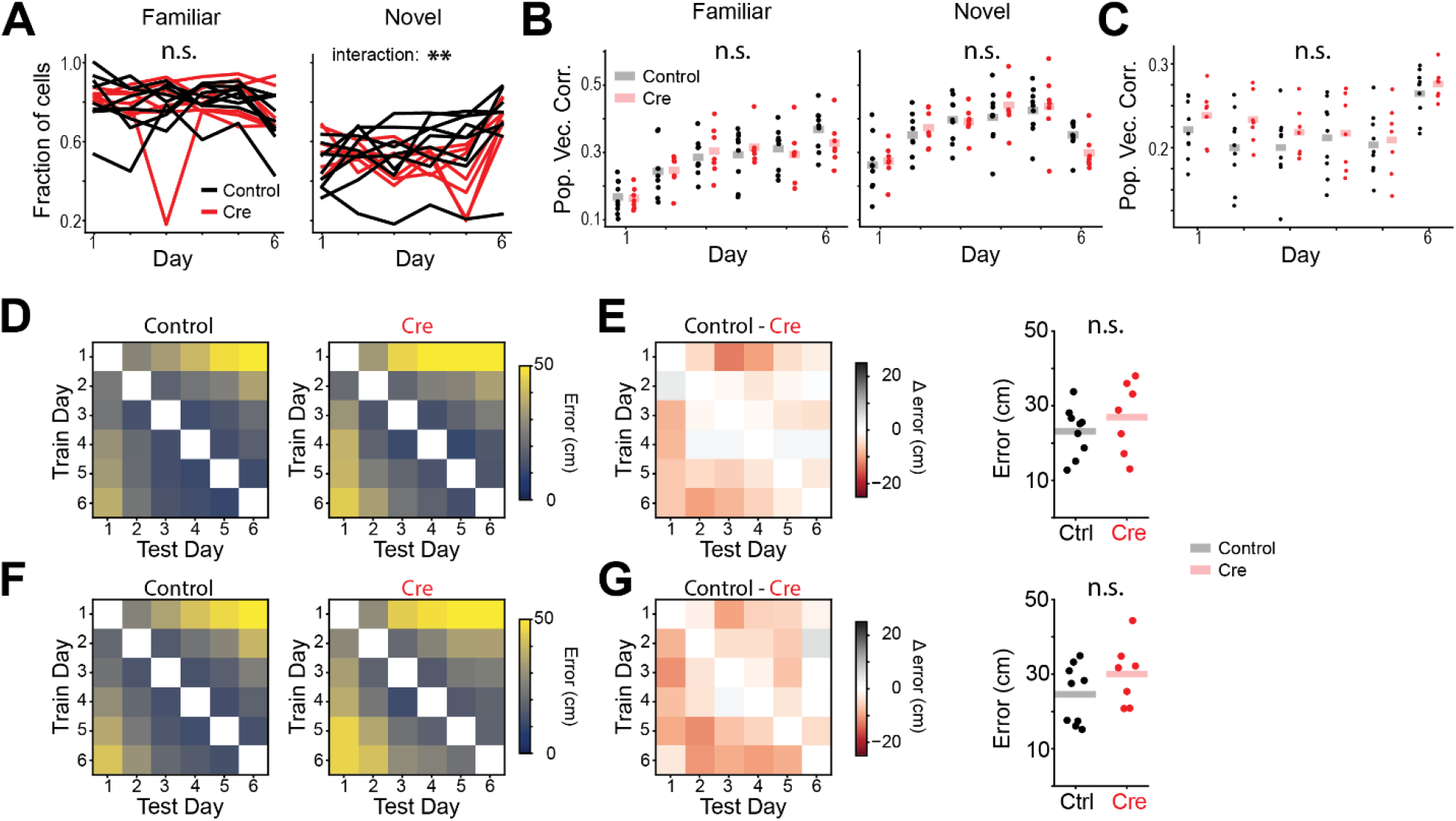
A) Fraction of cells with significant spatial information on each day for familiar (*Left*) and novel (*Right*) trials. Each line represents data from a single animal. We found a significant interaction between virus and day for novel trials but none of the posthoc tests were significant. [N=9 control mice, 7 Cre mice, 6 days. Familiar trials mixed effects ANOVA: virus main effect F(1,14)=.016 p=.795, day main effect F(5,70)=2.17 p=.067, interaction F(5,70)=.919 p=.474. Novel trials mixed effects ANOVA: virus main effect F(1,14)=.002 p=.966, day main effect F(5,70)=11.9 p=2.33×10-8, interaction F(5,70)=3.33 p=.009, all posthoc tests for control vs Cre difference on each day p>.05] B) Average trial x trial population vector correlation for familiar (*Left*) and novel (*Right*) trials on each day shows similar within-day stability between conditions. Each dot is the average for a single animal. [N=9 control, 7 Cre mice, 6 days. Familiar trials mixed effects ANOVA: virus main effect F(1,14)=.016 p=0.902, day main effect F(5,70)=34.1 p=1.69×10-17, interaction F(5,70)=1.15 p=.344. Novel trials mixed effects ANOVA: virus main effect F(1,14)=.016 p=.902, day main effect F(5,70)=43.8 p=3.15×10^−20^, interaction F(5,70)=2.76 p=.025] C) Population vector correlation between the average familiar trial activity and the average novel trial activity on each day shows comparable remapping across groups. [N=9 control, 7 Cre animals, 6 days. Mixed effects ANOVA: virus main effect F(1,14)=1.17 p=.298, day main effect F(5,70)=12.5 p=1.06×10^−8^, interaction F(5,70)=.586 p=.711] D-G) Across day decoding did not significantly differ between groups. We identified cells that we reliably tracked over all six days of the experiment. Naive Bayesian position decoding models were fit using the neural activity from one day and then tested on the other days. D) Across day decoding for familiar trials form control (*Left*) and Cre (*Right*) animals. E) *Left* - The difference (control-Cre) in across day decoding for familiar trials. *Right* - Average decoding error across days (average of matrix on the left for each mouse). [N=9 control mice, 7 Cre mice. Unpaired t-test, t=-1.32 p=.208] F) Same as (D) for novel trials. G) Same as (E) for novel trials. [N=9 control mice, 7 Cre mice, Unpaired t-test t=-.936 p=.365]

**Figure S4. Related to Figure 2.**
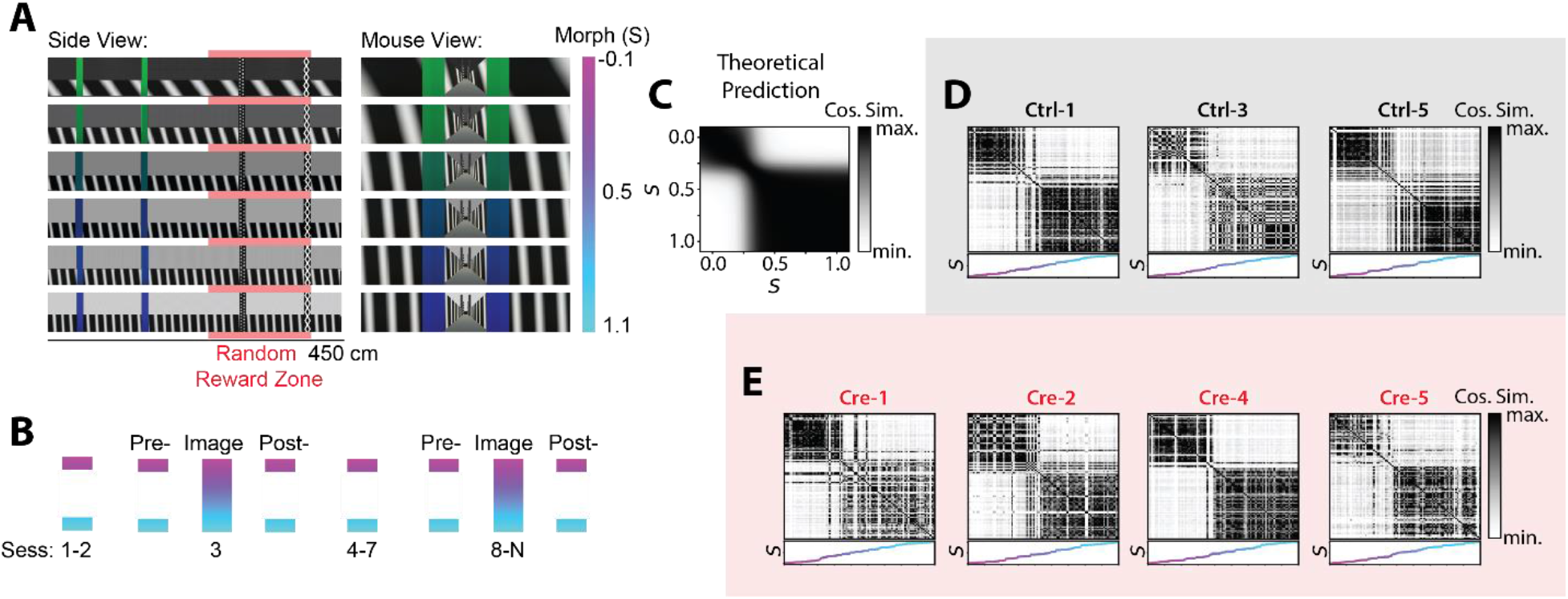
A) A subset of mice were trained in a separate VR task after the Y-maze task. Animals were trained to collect randomly placed rewards in a set of tracks that could be parametrically morphed between two extremes (see (71)). Example VR tracks that are gradually “morphed” between two extremes are shown (S-morphing parameter; side view - *Left*, mouse view - *Right*). Rewards were placed at random locations on the back half of the track on each trial (red highlighted region). B) Training protocol for inducing discrete remapping to intermediate morph values. Animals experienced randomly interleaved tracks of the extreme morph values except during a subset of imaging sessions (days 3 and 8-N). During these imaging sessions, morph values from the full range of stimuli were randomly interleaved. C) Trial x trial cosine similarity matrix for a simulated neural population that performs Bayesian inference to estimate which environment the animal is occupying. After training animals in the two extremes of the morph axis, the hippocampus should effectively discretize the stimuli into low and high morph values. The neural responses to low morph values should be highly similar, and the neural responses to high morph values should be highly similar. These two representations should have low similarity to one another. D) Example trial x trial population cosine similarity matrices for session 8 for example control mice. Note similarity to (C). E) Example trial x trial population cosine similarity matrices for session 8 for example Cre mouse. Note similarity to (C), indicating that more nuanced context discrimination dynamics are intact in animals lacking CA1 Stx3. Panels A & B reproduced from *(71)*.

**Figure S5. Related to Figure 3.**
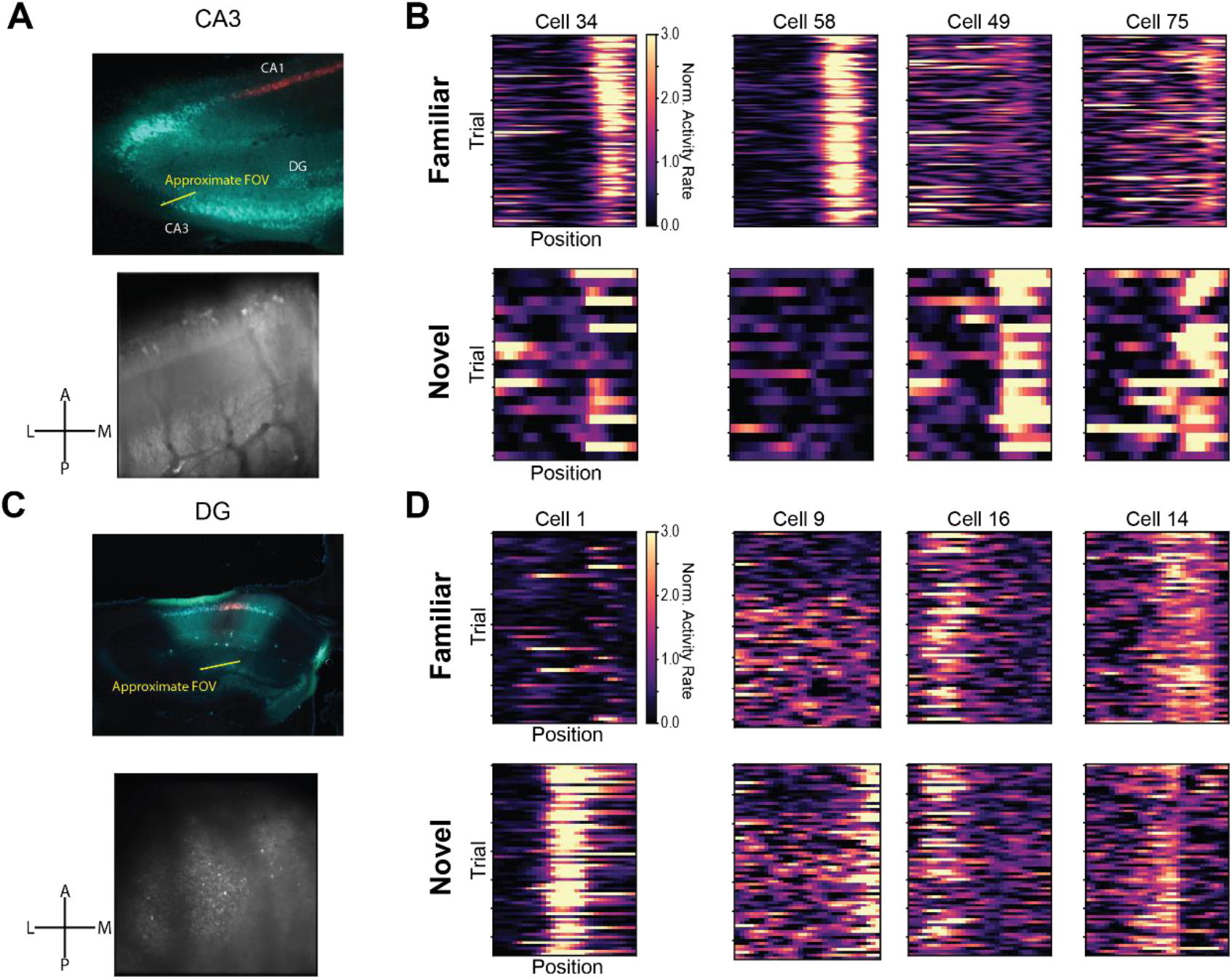
A) *Top* - histology from a mouse in which we were able to image CA2/3 place cells (Cre-8,DAPI-blue, GCaMP-green, mCherry - red). *Bottom* - the average two photon field of view from an example imaging session. Note the visible ventricle on the anterolateral edge of the 2-photon FOV. Crosshairs denote anterior-posterior (A-P) and lateral-medial (L-M) axes in FOV. B) Example trial x position activity rate maps from example CA2/3 place cells on day 1 for familiar (*Top*) and novel (*Bottom*) trials. Each column of plots indicates a different cell. Note stable place fields in both novel and familiar tracks. Most cells also remap. C) *Top* - histology from a mouse in which we were able to image dentate gyrus (DG) place cells through an intact CA1 (Cre-7, DAPI-blue, GCaMP-green, mCherry-red). *Bottom* - the average two photon field of view. We were able to image DG by focusing ∼600 microns past the CA1 imaging FOV. D) Example trial x position activity rate maps for DG place cells on an example session. This session was performed after all other training and was a repeat of the procedure from day 6 (Fig 1E). Note stable place cells in both novel and familiar trials. Cells also remap on the arms of the track.

**Figure S6. Related to Figure 4.**
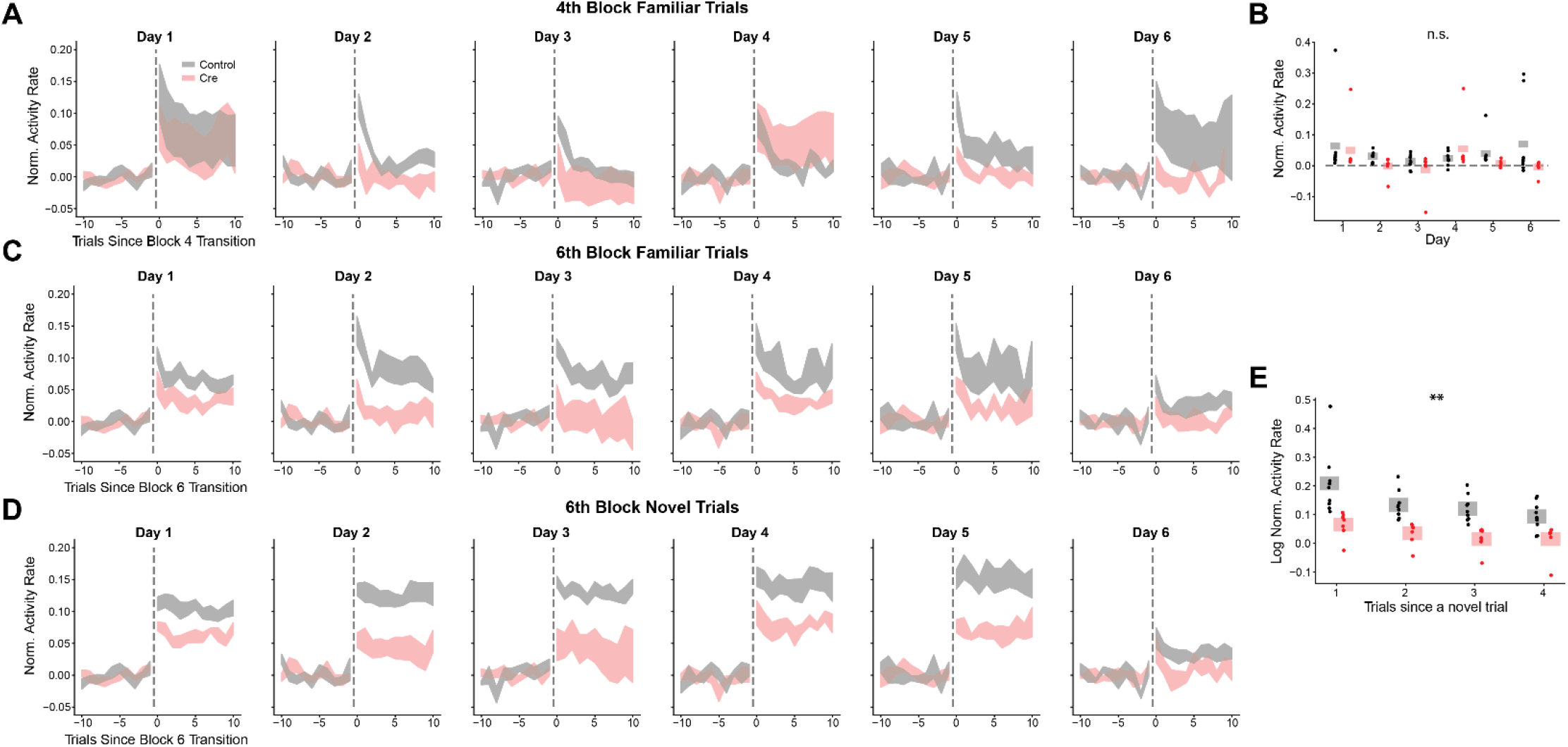
A) Population averaged normalized activity rate for familiar trials in the fourth trial block on each day as in Fig. 4C. Shaded region indicates mouse average ± SEM. In control and Cre animals, we occasionally observe a transient increase in activity rate at the start of this familiar block of trials. B) Averaged normalized activity rate on familiar trials in the fourth trial block for each mouse on each day. Dots indicate average normalized activity rate for each mouse. Shaded bars indicate across animal means. [N=9 control mice, 7 Cre mice, 5 days (day 6 excluded). Mixed effects ANOVA: virus main effect F(1,14)=.139 p=.715, day main effect F(4,56)=1.60 p=.187, interaction F(4,56)=.472 p=.756] C) Same as (A) for familiar arm trials in the last trial block on each day as in Fig. 4C. Day 1 and day 5 panels reproduced from Fig. 4C. D) Same as (A) for novel arm trials in the last trial block on each day as in Fig. 4C. Day 1 and day 5 panels reproduced from Fig. 4C. E) Familiar trial activity in the last trial block depends on time since encountering a novel trial. Across day averaged normalized activity rate in block 6 familiar trials plotted as a function of the number of trials since encountering a novel arm trial. Dots indicate the average activity rate for each mouse. Shaded bars indicate across animal mean. [N=9 control mice, 5 days, 4 trial conditions (# of trials since a novel trial). rmANOVA: day main effect F(5,40)=4.86 p=.001, trial since novel main effect F(2,16)=6.46 p=.009, interaction F(10,80)=1.44 p=.178; N=7 Cre mice, 5 days, 4 trial conditions (# of trials since a novel trial). rmANOVA: day main effect F(5,15)=2.50 p=.077, trial since novel main effect F(2,6)=15.2 p=.004, interaction F(10,30)=1.75 p=.116]

**Figure S7. Related to Figure 5.**
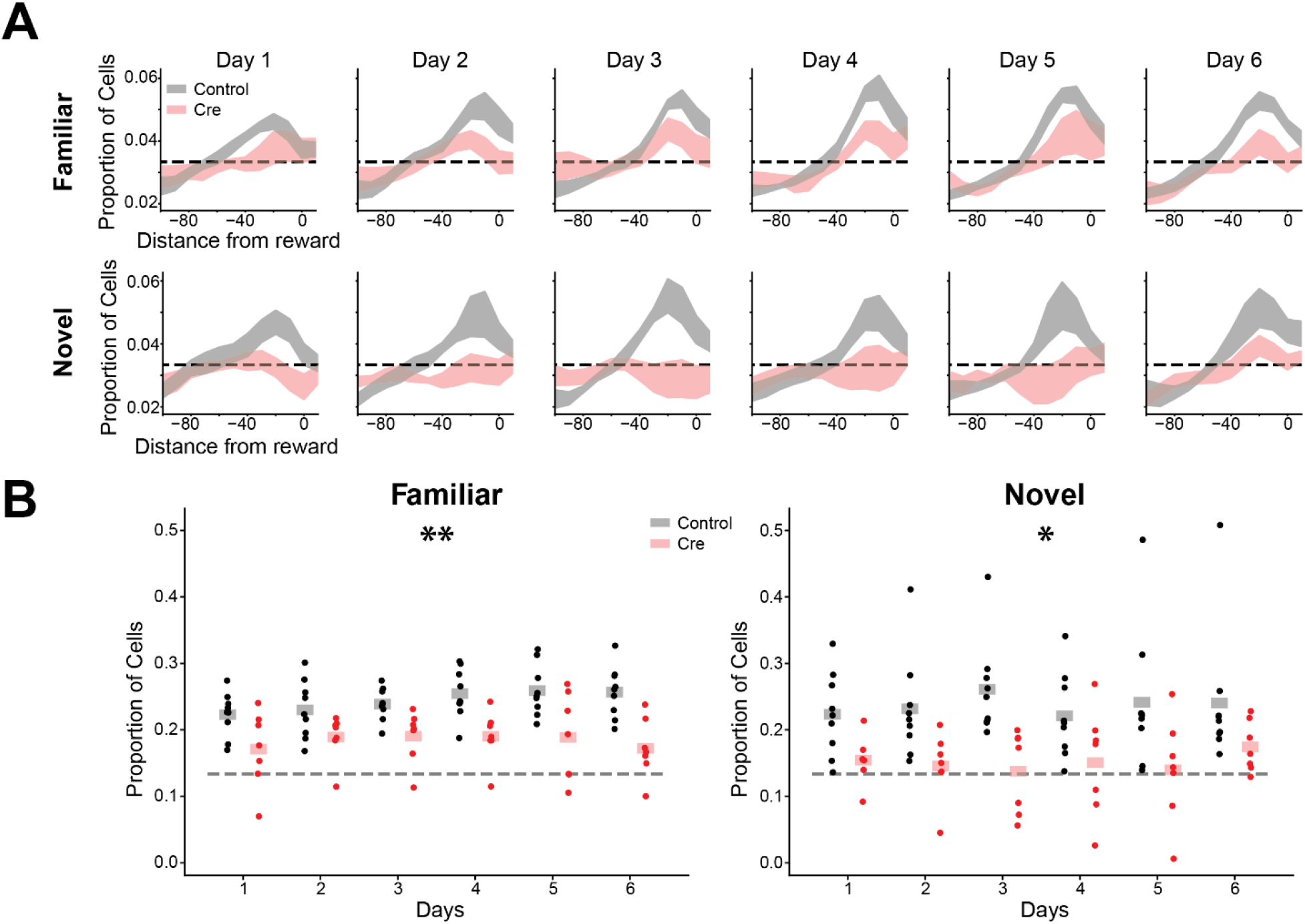
A) CA1 Stx3 is required for over-representation of reward locations. Proportion of place cells that have their peak in the peri-reward spatial bins for each day (*Top*-familiar, *Bottom*-novel). Data are plotted as across animal mean ± SEM (black-control, red-Cre). Dotted line indicates uniform distribution of place cell peaks across spatial bins. B) Proportion of place cells with their peak near the reward zone for each animal on each day (*Left* - familiar trials, *Right* - novel trials) [N=9 control mice, 7 Cre mice, 6 days. Familiar trials mixed effects ANOVA on logit transformed place cell proportions: N=9 control mice mice, 7 Cre mice, 6 days, virus main effect F(1,14)=9.45 p=.008, day main effect F(5,70)=1.10 p=.367, interaction F(5,70)=1.16 p=.337. Novel trials mixed effects ANOVA on logit transformed proportions: virus main effect F(1,14)=8.08 p=.013, day main effect F(5,70)=.908 p=.481, interaction F(5,70)=.947 p=.456]

**Figure S8. Related to Figure 6.**
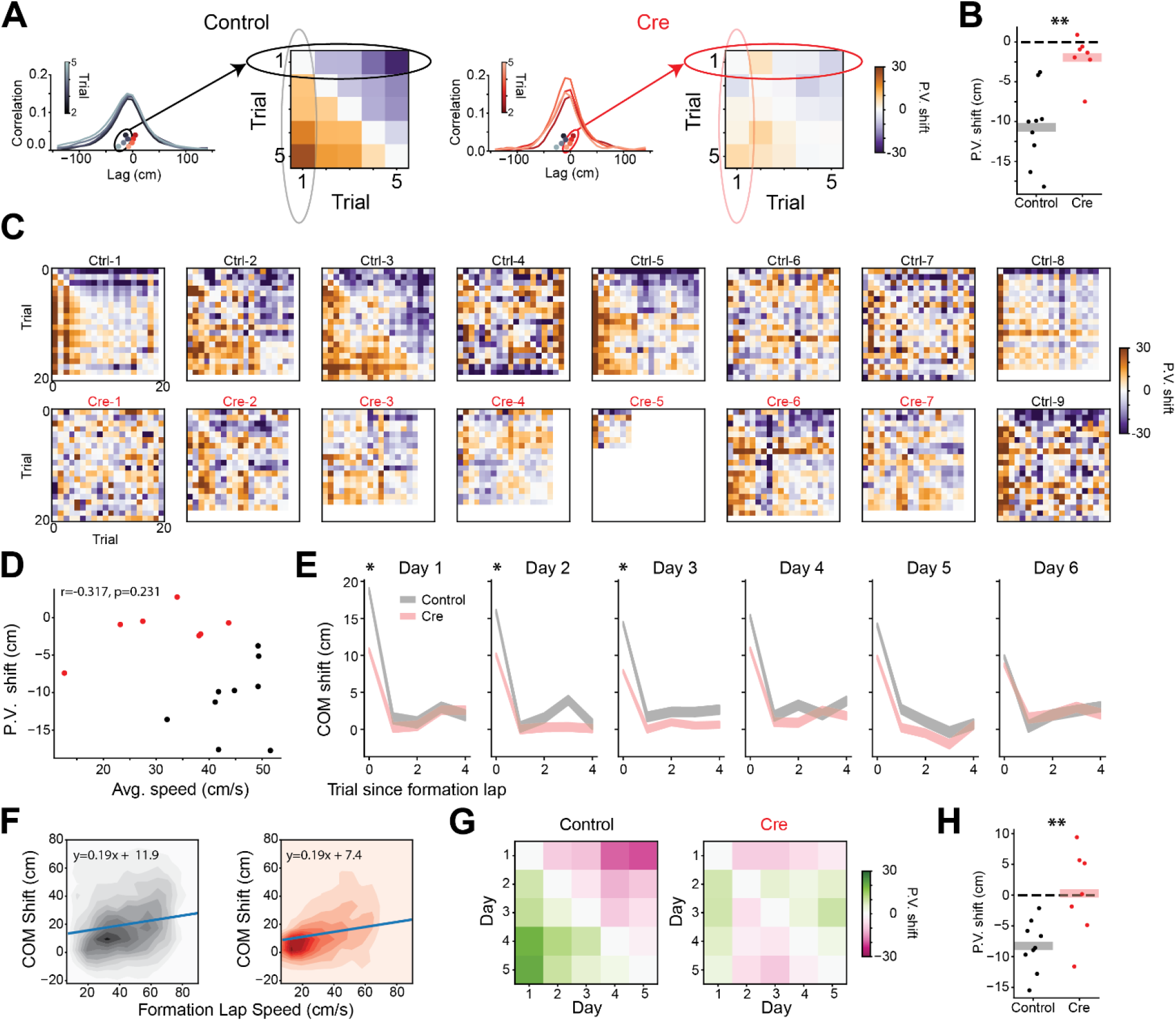
A) Schematic showing how the population vector (PV) shift is calculated. For each mouse we calculate the center of mass from trial x trial PV cross correlations. In the cross-correlation plots, reproduced from Fig 4D, the cross correlations of each trial with trial 1 are shown. These center of mass values yield a trial x trial antisymmetric PV shift matrix. The trial 1 examples give the entries in the first row/column of the matrix. The matrices shown are the across animal average PV shifts for each pair of trials. The average PV shift is calculated by averaging the upper triangle of this matrix. All data are from day 1 novel trials. B) Control mice display a greater backward PV shift than Cre mice. Replotted day 1 PV shift from Fig 4E. Dots indicate the PV shift for each mouse on day 1 novel trials. Shaded bars indicated across animal mean. [Control vs. Cre unpaired posthoc t-test: t=-4.63 p=.002 (Holm-corrected). Control one-sample t-test for significant shift from 0: t=-6.66 p=.002 (Holm-corrected). Cre one-sample t-test for significant shift from 0: t=-1.93 p=.710 (Holm-corrected)] C) PV shift matrices for all novel arm trials on day 1 for all mice. White space indicates trials that were not completed due to slower running speeds or stopping. Note that PV shift statistics in Fig 6 are calculated using only the first 5 trials. D) There is no significant correlation between PV shift and running speed in the first 5 novel trials of day 1. [N=9 control mice, 7 Cre mice. Pearson correlation: r=-.317 p=.231] E) The COM of place cell activity on each novel trial relative to the average place field COM, following the place field’s formation lap. Shaded regions indicate across place field mean +/− SEM (gray-control, red-Cre). After controlling for formation lap running speed through the place field, formation lap COM shifts were significantly greater on days 1, 2 and 3 for control animals. [Linear mixed effects regression. Wald test for significant mean effect of virus on trial 1 COM shift. Day 1: *χ*^2^ =6.37, p=.016, Day 2: *χ*^2^ =4.43, p=.035, Day 3: *χ*^2^ =6.07, p=.014] F) Trial 1 COM shift on novel trials on day 1 plotted against formation lap speed. Heatmaps show the joint histogram of COM shifts and place field formation lap speed through the place field (*Left* - Control, *Right* - Cre). Blue line shows the best fit regression for each condition from mixed effects linear regression (COM ∼ speed + virus). There is a difference in intercept values indicating a larger COM shift for control animals (see (E) for results of Wald test). G-H) Control but not Cre animals display a gradual backward shifting of place codes in novel trials across days. G) Across animal average across-day PV shift matrices, similar to (A), calculated using the trial-averaged novel arm activity in each mouse on each day. Instead of showing population shifts across trials, this analysis shows population shifts across days. H) Average across-day PV shift for each mouse. Dots represent the average of the upper triangle of the PV shift matrix as in (G) for each mouse. Shaded bars indicate across mouse average. [N=9 control, 7 Cre mice, unpaired t-test: t=-3.00, p=.010. Control one-sample t-test for significant shift from 0: t=-5.95 p=3.42×10-4. Cre one-sample t-test for significant shift from 0: t=.111 p=.915]

**Figure S9. Related to Figure 7.**
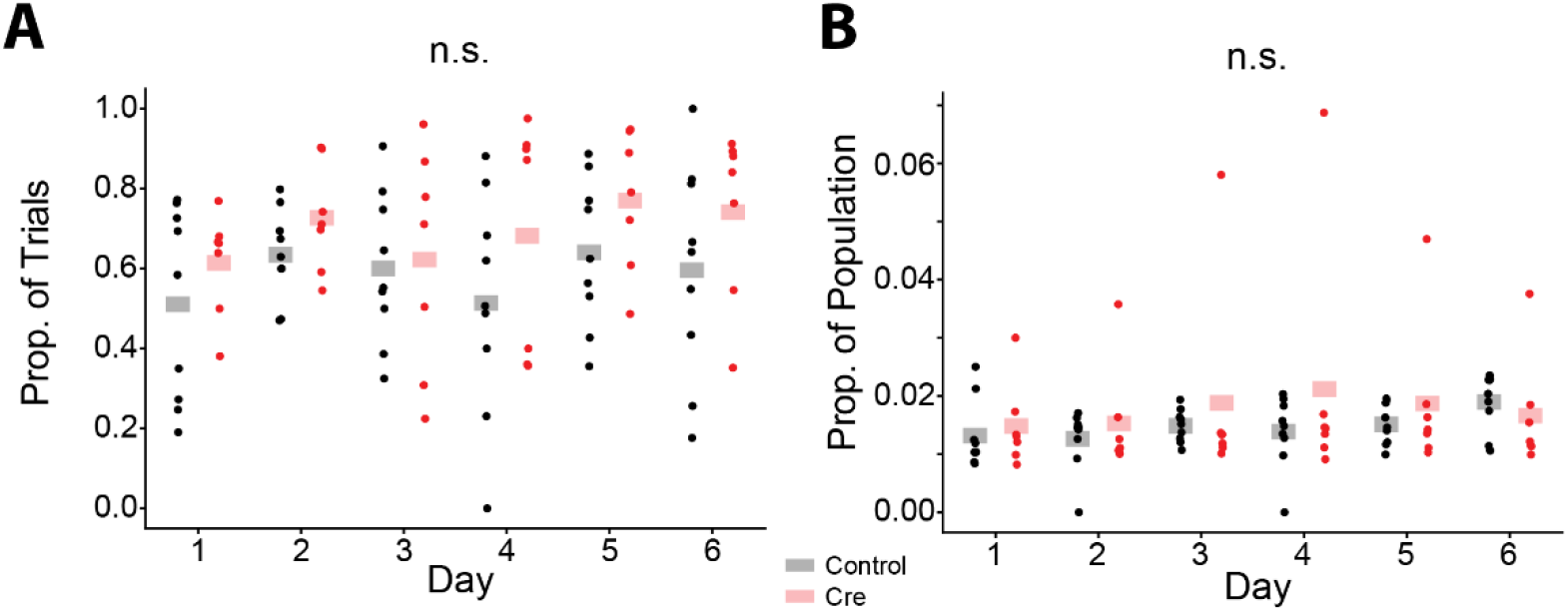
A) Proportion of trials that contain a significant HSE does not differ between control animals (black) and Cre animals (red). Individual dots represent the proportion of trials on a particular day for each mouse. Shaded bars indicate the across animal mean on each day. [N=9 control, 7 Cre mice, 6 days. Mixed effects ANOVA: virus main effect F(1,14)=2.13 p=.167, day main effect F(5,70)=1.32 p=.364, interaction F(5,70)=.383 p=.859] B) Proportion of the neural population that participates in HSEs does not differ between control (black) and Cre (red) animals. Each dot represents the average size of HSEs for each mouse quantified as the proportion of the population participating. Shaded bars indicate across animal mean on each day. [N=9 control, 7 Cre mice, 6 days. Mixed effects ANOVA: virus main effect F(1,14)=.397 p=.539, day main effect F(5,70)=2.30 p=.054, interaction F(5,70)=1.88 p=.108]

## Methods

### Subjects

All procedures were approved by the Institutional Animal Care and Use Committee at Stanford University School of Medicine. For all hippocampal experiments, we used male *Stx3^fl/fl^* mice^135^ due to the known effects of estrous cycle on hippocampal LTP. All mice were housed in groups of between two and five same-sex littermates. After surgical implantation or virus injection, mice were housed in transparent cages and kept on a 12-hour light/dark schedule, and mice allocated to head-fixed behavior were housed with a running wheel from ∼1 month of age. All experiments were conducted during the light phase. Mice were between 2-4 months at the time of virus injection for freely moving behavior experiments and 3-5 months at the time of surgery for two photon experiments.

### Statistics

All numerical data in figures are means ± SEM. Freely moving behavior experiments were done in a blind manner. Statistics on the freely moving behavior were run using Prism 9 (GraphPad). Analyses of virtual reality behavior and Ca^2+^ imaging data: Generalized linear mixed effects models were run using the statsmodels package (https://www.statsmodels.org/stable/index.html). Mixed effects ANOVAs and subsequent posthoc t-tests were run using the pingouin package (https://pingouin-stats.org/) with a Holm step-down Bonferonni correction for multiple comparisons. No statistical methods were used to pre-determine the number of mice to include in this study, but our sample sizes are similar to those reported in previous publications.

### Stereotaxic injection for freely-moving behaviors

AAVs (serotype DJ, produced by HHMI Janelia Resarch Campus) expressing Synapsin-EGFP-CRE and Synapsin-EGFP-ΔCre were injected into *Stx3^fl/fl^* mice. Mice were anesthetized with tribromoethanol (250 mg/kg, both viral conditions for open field and elevated plus maze) or ketamine (100 mg/kg)/xylazine (10 mg/kg) (T-Maze) and head-fixed with a stereotaxic device. A glass micropipette with a long, narrow tip was pulled using a micropipette puller to deliver the virus. The glass pipette was slowly lowered to the target area and left for 2 min before virus injection. Virus solution was injected at an infusion rate of 100 nl/min and withdrawn 5 min after the end of injection. 300 nL of virus was infused into 4 sites – bilaterally in 2 different anterior-posterior positions (AP: −2 and −2.5, ML: 1.42, DV: −1.24). The mice were used for experiments at least 2 weeks after the virus infusion.

### Behavior

All mice were handled for at least 3 days (>5 min/day) before behavioral experiments. All video tracking experiments except fear conditioning were analyzed with the Viewer III tracking system (Biobserve) followed by custom MATLAB scripts.

#### Open field

Mice were individually placed into the center of the open field chambers (34cm × 34cm × 40 cm, white) and allowed to freely explore for 30 min. The total traveling distance and time spent in the center area were analyzed.

#### Elevated plus maze

The gray maze had four 30cm × 8 cm arms. Two “open” arms did not have walls, while 10 cm high walls enclosed the two “closed” arms. The maze was elevated 40 cm over the floor. At the beginning of the test, each mouse was individually placed at the junction of an open and a closed arm, facing the open arm, and was allowed to freely explore for 5 min. The time spent in the open versus the closed arms was analyzed.

#### Freely-Moving Novelty T-Maze

Adapted from ref ^54^. The dimensions of each transparent arm were 30cm×10cm×20cm (L×W×H). During training, the entrance to one arm (“Novel arm”, assignment counterbalanced across groups) was blocked with black, matte plastic. Training consisted of five, 2min sessions separated by a 1 minute ITI in which each subject was allowed to explore the start and “familiar” arms. Following an ITI, each mouse was allowed to explore all arms of the maze – the novelty preference test. This session was 2min and began when all 4 paws left the start arm. Test performance was measured by the difference score: (time spent in the novel arm) – (time spent in the familiar arm).

### Histology/Imaging

Mice were deeply anesthetized with tribromoethanol and transcardially perfused with PBS followed by 4% PFA. Brains were removed and post-fixed in 4% PFA for 24 hours (1 hour if proceeding with X-gal staining) and then transferred to 30% sucrose in PBS for 48 hours at 4°C. 100 μm sections were taken on a cryostat (CM3050 S; Leica Biosystems) and mounted on glass slides with DAPI Fluoromount-G (Southern Biotech). Images were collected using a 10x objective on a VS120 or VS200 slide-scanning microscope (Olympus). We annotated standard Paxinos plates with viral spread from each mouse we used in freely-moving behavior.

#### X-gal staining

Slides with sections were rinsed in PBS 3 times, each for 5 min. Sections were then developed in a solution containing 5 mm K-ferricyanide, 5 mm K-ferrocyanide, 2mM MgCl_2_ and 1 mg/ml X-gal) at 37°C for 24 h in the dark. Finally, samples were rinsed with distilled water twice, slightly air-dried, and mounted with coverslips. Transmitted light imaging was then performed.

### *In vivo* imaging virus injections and window implants

*Stx3^fl/fl^* mice were first anaesthetized by an intra-peritoneal injection of a ketamine/xylazine mixture (∼85 mg/kg). Animals were also subcutaneously administered 0.08 mg Dexamethasone and 0.2 mg Carprofen. After the first hour of the procedure, anesthesia was maintained with 0.5-2% isofluorane. For control mice (n=9 Ctrl1-9), adeno-associated virus containing GCaMP7f (AAV1-hSyn-jGCaMP7f, AddGene viral prep #104488-AAV1) and a second virus encoding the static red indicator mCherry (AAVDJ-hSyn-mCherry, Stanford Gene Vector and Virus Core) were mixed and drawn into the same 36 gauge Hamilton syringe (World Precision Instruments; GCaMP titer - 1 x 10^12^, mCherry titer - 5 x 10^12^). For cre-injected mice (n=7, Cre1-7), the same GCaMP virus and a virus encoding Cre recombinase and mCherry (AAVDJ-CaMKII-mCherry-IRES-cre, Stanford Gene Vector and Virus Core) were mixed and drawn into the syringe (cre titer – 5-10 x 10^12^). For both groups of mice, virus was injected in two locations in the left dorsal CA1-region (injection 1: −2 AP, −1.4 ML, 1.35 DV; injection 2: −2.5 AP, −1.4 ML, 1.35 DV) and two locations in the right dorsal CA1-region (injection 1: −2 AP, −1.4 ML, 1.35 DV; injection 2: −2.5 AP, −1.4 ML, 1.35 DV). Each injection was 300 *η*L (50 *η*L/min) and the needle was left in place for 5 minutes following each injection to allow for virus dispersal. In an additional mouse (Cre-8), we performed identical dorsal CA1 injections of the cre virus without GCaMP and then injected the same GCaMP virus into the left dorsal CA3 (−1.5 AP, −1.6 ML, 1.98 DV, 500 *η*L at 50 *η*L/min).

To provide optical access to the CA1 pyramidal cell layer, after virus injections were complete, an imaging cannula was implanted over the left dorsal hippocampus as previously described^71,136^. Imaging cannulas consisted of a 1.3 mm length stainless steel cannula (3 mm outer diameter, McMaster) glued to a circular cover glass (Warner Instruments, #0 cover glass 3mm diameter; Norland Optics #81 adhesive). Excess glass overhanging the edge of the cannula was shaved off using a diamond tip file. A 3 mm diameter craniotomy was performed over the left posterior cortex (Cre1-7 and Ctrl1-9 centered at −2 mm AP, −1.8 mm ML, Cre8 centered at −1.5 mm AP, −2 ML). The dura was then gently removed and the overlying cortex was aspirated using a blunt aspiration needle under constant irrigation with ice cold sterile artificial cerebrospinal fluid (ACSF). Excessive bleeding was controlled using gel foam that had been torn into small pieces and soaked in sterile ACSF. Aspiration ceased when the fibers of the external capsule were clearly visible. Once bleeding had stopped, the imaging cannula was lowered into the craniotomy until the coverglass made light contact with the fibers of the external capsule. In order to make maximal contact with the hippocampus while minimizing distortion of the structure, the cannula was placed at approximately a 10 degree roll angle relative to the animal’s skull. The cannula was then held in place with cyanoacrylate adhesive. A thin layer of adhesive was also applied to the exposed skull. A number 11 scalpel was used to score the surface of the skull prior to the craniotomy so that the adhesive had a rougher surface on which to bind. A stainless steel headplate with a left offset 7 mm diameter beveled window was placed over the secured imaging cannula at a matching 10 degree angle and cemented in place with Met-a-bond dental acrylic that had been dyed black using India ink.

At the end of the procedure, animals were administered 1 mL of saline and .2 mg of Baytril and placed on a warming blanket to recover. Animals were typically active within 20 min and were allowed to recover for several hours before being placed back in their home cage. Mice were monitored for the next several days and given additional Carprofen and Baytril if they showed signs of discomfort or infection. Mice were allowed to recover for 14 days before beginning water restriction and VR training.

### Two photon (2P) imaging

To image the calcium activity of neurons, we used a resonant-galvo scanning two photon microscope (Neurolabware). 980 nm (Ctrl1-5, Cre1-7) or 920 nm (Ctrl6-9, Cre8) light (Coherent Discovery laser) was used for simultaneous stimulation of GCaMP and mCherry in CA1. 920 nm light was used in cases where mCherry fluorescence bleedthrough into the green photomultiplier tubes exceeded usable levels when stimulating at 980 nm. 920 nm light was also used to image CA2/3 and dentate gyrus (DG). Laser power was controlled using a pockels cell (Conoptics). Average power for excitation ranged from ∼35 mW to ∼80 mW measured at the front face of the objective (CA1 imaging: Nikon 16x, 3 mm WD, 0.8 NA; DG & CA3 imaging: Leica 25x, 3 mm WD, 1.0 NA). A 512 x 796 pixel field of view (FOV) was collected using unidirectional scanning at 15.46 Hz. Cells were imaged continuously under constant laser power during each block of trials (described below). To minimize photodamage and photobleaching, the pockels cell was used to reduce laser power to minimal levels between trial blocks.

Putative pyramidal cells were independently identified using the Suite2P software package (https://github.com/MouseLand/suite2p) on each session. Segmentations were curated by hand to remove ROIs that contained multiple somas, dendrites, or contained cells that did not display a visually obvious transient. Cells that did not contain a clear mCherry signal were also removed. The same FOV was imaged during each session. Custom code was used to find cells that were successfully tracked across imaging sessions. Baseline fluorescence was calculated for each cell within each block of trials independently using a sliding window maximin procedure (modification of default suite2p procedure). *Δ*F/F was then calculated for each cell as the change in fluorescence from baseline divided by baseline. These traces were median filtered to suppress optical shot noise and then deconvolved with a canonical calcium kernel to obtain “activity rates”.

### Virtual reality (VR) design

All virtual reality environments were designed and implemented using the Unity game engine (https://unity.com/). Virtual environments were displayed on three 24 inch LCD monitors that surrounded the mouse and were placed at 90 degree angles relative to each other. A dedicated PC was used to control the virtual environments and behavioral data. This PC was synchronized with calcium imaging acquisition using TTL pulses sent to the scanning computer on every VR frame. Mice ran on a fixed axis foam cylinder and running activity was monitored using a high precision rotary encoder (Yumo). Separate Arduino Unos were used to monitor the rotary encoder and control the reward delivery system.

### Water restriction and VR training

To incentivize mice to run, animals’ water intake was restricted. Water restriction was implemented 10-14 days after the imaging cannula implant procedure. Animals were given 0.8 - 1 mL of 5% sugar water each day until they reached ∼85% of their baseline weight and given enough water to maintain this weight.

Mice were handled for 3 days during initial water restriction and watered through a syringe by hand to acclimate them to the experimenter. On the fourth day, we began acclimating animals to head fixation (day 4: ∼30 minutes, day 5: ∼1 hour). After mice showed signs of being comfortable on the treadmill (walking forward and pausing to groom), we began to teach them to receive water from a “lickport”. The lickport consisted of a feeding tube (Kent Scientific) connected to a gravity fed water line with an in-line solenoid valve (Cole Palmer). The solenoid valve was controlled using a transistor circuit and an Arduino Uno. A wire was soldered to the feeding tube and capacitance of the feeding tube was sensed using an RC circuit and the Arduino capacitive sensing library. The metal headplate holder was grounded to the same capacitive-sensing circuit to improve signal to noise, and the capacitive sensor was calibrated to detect single licks. The water delivery system was calibrated to deliver ∼4 *μ*L of liquid per drop.

After mice were comfortable on the ball, we trained them to progressively run further distances on a VR training track to receive sugar water rewards. The training track was 450 cm long with black and white checkered walls. A pair of movable towers indicated the next reward location. At the beginning of training, this set of towers was placed 30 cm from the start of the track. If the mouse licked within 25 cm of the towers, it would receive a liquid reward. If the animal passed by the towers without licking, it would receive an automatic reward. After the reward was dispensed, the towers would move forward. If the mouse covered the distance from the start of the track (or the previous reward) to the current reward in under 20 seconds, the inter-reward distance would increase by 10 cm. If it took the animal longer than 30 seconds to cover the distance from the previous reward, the inter-reward distance would decrease by 10 cm. The minimum reward distance was set to 30 cm and the maximal reward distance was 450 cm. Once animals consistently ran 450 cm to get a reward within 20 seconds, the automatic reward was removed and mice had to lick within 25 cm of the reward towers in order to receive the reward. After the animals consistently requested rewards with licking, we began novel arm Y-maze training described below. Training took 2 - 4 weeks.

### VR novel arm Y-maze

We sought to create a VR task that could probe the same phenomena as the freely moving T-maze behaviors. However, by design, these tasks result in dramatic occupancy differences between the arms of the maze. This complicates statistical analyses of spatial and contextual coding. Furthermore, it is unknown if wildtype mice prefer to explore novel VR environments. To deal with this tradeoff we created a virtual Y-maze in which we forced mice to turn down different arms of the maze (Fig 1E, Fig S2A, [Movie S1](https://www.dropbox.com/scl/fi/hyru8d90p7rfoqwbhh3uo/Supplementary-Video1-1080p.mp4?rlkey=q14fq3tbk0m5komaflodhweno&dl=0)). Briefly, on each trial, the mouse began in a dark “timeout box” at the beginning of the track. The mouse self-initiated the trial by running forward on the ball. The animal then ran down the track to find a hidden reward and ran to the end of the track to end the trial. Whether the trial was a left or right turn was predetermined. The mouse was then teleported back to the “timeout box”, where it was forced to wait for a brief random duration (1-5 seconds, uniformly distributed) before beginning the next trial. This design did not allow us to assess the animal’s preference for each arm; however, it allowed us to test the animal’s ability to distinguish the arms and examine novel context encoding while controlling for the occupancy of the different arms. The task essentially reduces to previously published VR novel context tasks^23,26^ while avoiding the sudden change in stimuli used in these tasks.

The animal needed to run 300 cm to complete each trial, but apparent motion along the track was a nonlinear function of movement on the ball. Smooth trajectories for left and right turns were created by fitting a cubic spline to a set of control points along the track. The left and right turn control points were symmetric, so that left and right turns were mirrored. The stem and arms of the maze were of equal length, but to create a smooth trajectory, the trajectories noticeably diverge approximately two thirds of the way down the stem of the Y-maze. Due to the uneven spacing of control points to create convincing trajectories, the visual speed as a function of movement on the ball slows near the turn and increases at the ends of the maze. As the spline trajectory parameter is linearly related to the animal’s motor action, we perform all spatially binned analyses as binned values of the spline parameter. These bins correspond to 10 cm of movement along the ball (30 bins to cover the track).

After initial running training, animals were trained to run in a “training Y-maze” to acclimate them to the spline trajectories. All walls of the training Y-maze were black and white checkerboards, with a black floor and a gray ceiling. The mice were required to lick to receive a hidden reward (i.e. no explicit visual cue marked the location of the reward) at the midpoint of either arm of the maze. Left and right turns were randomly interleaved on the training Y-maze. Once the animals ran smoothly and licked consistently to get the hidden rewards (1-3 days), we began the novel arm Y-maze task.

For the novel arm Y-maze, each section of the track had a dramatically different pattern of visual stimuli to aid the mice in distinguishing the arms (Fig S2A, [Movie S1](https://www.dropbox.com/scl/fi/hyru8d90p7rfoqwbhh3uo/Supplementary-Video1-1080p.mp4?rlkey=q14fq3tbk0m5komaflodhweno&dl=0)). The reward was omitted on 10% of trials. On each of the first 5 days of the experiment, the animals performed 6 blocks of trials. For the first 5 blocks of trials, animals were forced to go down only one arm (“familiar” arm) and the view of the other arm was blocked with a translucent virtual curtain (Fig S2A, [Movie S1](https://www.dropbox.com/scl/fi/hyru8d90p7rfoqwbhh3uo/Supplementary-Video1-1080p.mp4?rlkey=q14fq3tbk0m5komaflodhweno&dl=0)). In the sixth block, the virtual curtain was removed and familiar and novel arm trials were randomly interleaved. On the sixth day, the block structure was retained but familiar and novel trials were randomly interleaved from the first trial. Mice were randomly assigned to having the left or right arm as the familiar arm. To determine if the mice could distinguish the arms of the maze, they were required to lick at a hidden zone (25 cm wide) at the beginning of the left arm and the end of the right arm in order to receive a liquid reward. On the first block of trials on day 1, an automatic reward was dispensed at the end of the reward zone if the mouse did not lick within the reward zone. To mimic the freely moving task while getting sufficient trials to perform statistical comparisons, each of the first 5 blocks was a maximum of 5 minutes or 20 trials and the last block was 10 minutes or 40 trials. The inter-block interval was 1 minute.

On day 7, the procedure from day 6 was repeated for the first 2 blocks of trials. During the inter-block interval, we switched the reward locations on the left and right arms without changing any of the visual stimuli. An automatic reward was available on block 3 but removed for the remaining blocks. On day 8, the rewards remained in the reversed locations and left and right trials were randomly interleaved for all blocks.

### VR behavioral analyses

Anticipatory lick rate was calculated as the average lick rate (licks/sec) in the 30 cm preceding the reward zone. To normalize for differences in overall lick rate, lick rate in each spatial bin was divided by the overall average lick rate for each animal. Similarly, normalized running speed was calculated as running speed in each spatial bin was divided by the animal’s average running speed.

To analyze slowing in response to novelty, average running speed in the 6th block of trials was divided by the average running speed in the 10 trials preceding the start of the 6th block. We took the *log*_10_ of the resulting value so that decreases in running speed are negative values and increases in running speed are positive values.

Coefficient of variation (CV) of running speed was calculated as the standard deviation of running speed across trials in each spatial bin divided by the mean running speed across trials in each spatial bin. Speed CV values were compared between groups in the 30 cm preceding the reward zone.

### Place cell identification and analyses

Place cells were identified independently on familiar and novel trials on each day using a previously published spatial information (SI)^137^ metric 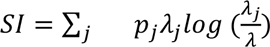. Where *λ*_*j*_ is the average activity rate of a cell in position bin *j*, *λ* is the position-averaged activity rate of the cell and *p*_*j*_ is the fractional occupancy of bin *j*.

To determine the significance of the SI value for a given cell, we created a null distribution for each cell independently using a shuffling procedure. On each shuffling iteration, we circularly permuted the time series of the cell relative to the position trace within each trial and recalculated the SI for the shuffled data. Shuffling was performed 1,000 times for each cell, and only cells that exceeded 95% of permutations were determined to be significant place cells.

Place cell sequences (Fig 2E-G & 5A) were calculated using a split halves procedure. The average firing rate maps from odd-numbered trials were used to identify the locations of peak activity and the activity on even-numbered trials is shown (z-scored across positions). When comparing place cell sequences for different trial types, we considered the union of place cells in both conditions.

### Single place field analyses

Significant place fields were identified by a separate method independent of place cell identification. To identify candidate significant place fields, we first calculate the trial averaged activity rate map for each cell. We then created a null distribution for these rate maps by circularly permuting the time series of the cell relative to the position trace within each trial each individual trial’s rate map 1,000 times. Contiguous spatial bins with an activity rate that exceeded 95% of this null distribution were considered potential place fields. Potential fields that were smaller than 20 cm were removed.

After these potential place fields were identified, we implemented additional filtering similar to previous publications^61^. We first identified the place field formation lap. A trial was considered the formation lap if it was the first trial in which the cell displayed significant calcium activity in the place field and then continued to display significant calcium activity in at least 66% percent of the remaining trials (minimum 4 remaining trials). Significant calcium activity was defined as the deconvolved activity rate exceeding 20% of the cell’s maximum activity rate in the place field. Potential place fields in which we could not identify a formation lap were removed from further single place field analyses.

The center of mass (COM) of place field firing on each lap was calculated using the non-spatially binned running speed and calcium activity to retain maximal resolution.

### Naïve Bayes decoding

To decode position from spatially binned population neural activity, we used the following decoder: 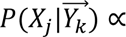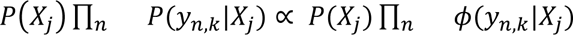, where 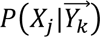 is the posterior probability that the mouse is in position *X_j_* given the position binned vector of neural activity rates 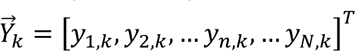 where *n* indexes the neuron identity and *k* indexes the unknown position. *P*(*X_j_*) is the prior probability the mouse is in position bin 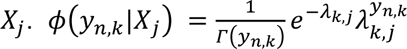, is a continuous quasi-Poisson potential function, where *λ_k,j_* is the trial-averaged activity rate of neuron *k* at position *j*. We assume that *P*(*X*) is uniform, reducing the model to a maximum likelihood decoder.

We used leave-one-trial out cross validation to assess the accuracy of the decoder. Error was calculated as the mean absolute deviation of the most likely position, 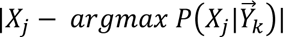. To compare errors across animals, we repeatedly randomly sampled an equal number of cells from each mouse with replacement to perform cross-validation (50 iterations per mouse). For decoding across days, all cells that were tracked across the first 6 days of the experiments were used in the decoder.

### K-Winners-Take-All (KWTA) model for place cell inheritance

Our goal for this model was to write down the simplest network with plausible components that could simulate whether a CA1 population could inherit a place cell representation even without synaptic plasticity and, if so, how plasticity could change this representation. This model has only three main components: 1) spatially selective input neurons, 2) Hebbian plasticity between input model CA3 and CA1 neurons and, 3) recurrent inhibition between model CA1 neurons.

We considered a two layer neural network with a set of spatially selective input neurons, 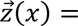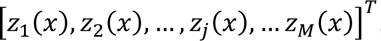, a set of output neurons, 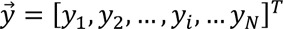, and a connectivity matrix, *W*, where *W_ij_* is the weight from input neuron *j* to output neuron *i*. Input neurons have radial basis function tuning for position, 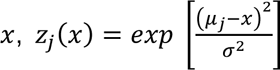, where *μ*_j_ is the center of the radial basis function for neuron *j*. Basis functions were chosen to tile positions across the population, [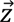. *σ*^2^ is the width of the radial basis function and is fixed for all input neurons. For Fig. 3, M=1000, N=1000, *σ*^2^ = 0.5, *x* ∈ [0,10].

We accomplished recurrent inhibition between output neurons using a K-Winners-Take-All approach. At every position along the track *y_i_* = {*KWTA*(*a*(*x*)) ⊙ *a*(*x*) + *σ_y_*, 0}, where 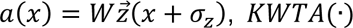 is a vector-valued function that applies the KWTA threshold and outputs a binary vector choosing the K winners, ⊙ denotes the elementwise product and *σ*_*y*_ is additive noise to the output. *σ_z_* is a stimulus noise term. K=50, *σ_y_*∼0.5*N*(0,1), *σ_z_*∼0.05*N*(0,1).

Weights were updated according a basic Hebbian learning rule at the end of each track traversal, *ΔW_ij_* = *ηz_j_y_i_*, where *η* is a constant. We required that weights had to be nonnegative. Weights also decayed to prevent explosive growth of synapse weight known to occur in Hebbian models, and additive Gaussian noise was included in the weight update. This gave the following weight update equation: 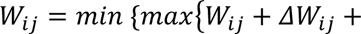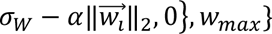. For the “no Plasticity” model *η* = 0, *α* = 0 and for the “Plasticity” model *η* = 1 *x* 10^−4^, *α* = 9*x*10^−4^. For all simulations *σ*_*W*_ = 0.001*N*(0,1) and *w_max_* = 10.

We initialized weights using a lognormal distribution (mean=0, standard deviation = 0.5)^138,139^ though uniform weights also reproduced our results under different noise regimes (data not shown).

### Novel arm population activity rate

To assess whether overall neural activity rate increased in a block of trials while controlling for bleaching or within session FOV drift, each cell’s position-binned activity rate was divided by the mean activity rate of that cell in the 10 trials preceding the block of trials to be analyzed. We took the *log*_10_ of the mean of this normalized activity rate across all cells to get a population normalized activity rate. This normalized value is less than zero when the population decreases its activity relative to baseline, and it is greater than zero when the population increases activity relative to baseline.

### Reward overrepresentation analysis

Place cells were considered to be near the reward zone if their location of peak activity was in the 50 cm preceding the front of the reward zone. To find “reward cells”, we analyzed only neurons that had significant spatial information in both familiar and novel trials. If the neuron’s location of peak activity was within 50 cm of the front of the reward zone in both trial types, it was considered a “reward cell”

### Population vector cross-correlation

Using a method previously described^60^, we estimated trial x trial spatial shifts in population activity. For each pair of trials, we shifted one trial relative to the other (±300 cm, 10 cm spatial bins) and computed the population vector correlation at each lag (i.e. a spatial cross-correlation). We calculated the shift in population activity as the center of mass (COM) of this cross-correlation (Fig 6A-C & Fig S7). This gave an anti-symmetric matrix of COM values. To calculate the average initial shift of the population we averaged the upper triangular entries in the trial x trial matrix containing the first five novel arm trials.

### Highly Synchronous Events (HSEs)

HSEs were identified using a novel shuffling procedure. We first identified periods following reward delivery where the animals were stationary (running speed <2 cm/s) for at least 750 msec. Next, we identified cells that displayed a significant calcium transient that began after the animal stopped and the reward was delivered. A significant calcium transient was considered as activity that exceeded 20% of the cell’s maximum activity rate in that session. Cells were considered coactive if their significant transients began within a 500 msec window.

To determine the number of cells that were expected to be coactive by chance in each imaging session, we circularly permuted each cell’s timeseries independently within each eligible stopping period and recalculated the number of coactive cells at each timepoint. This procedure was repeated 1,000 times, and the threshold for a significant HSE was chosen as the 95th percentile of this shuffled distribution.

